# A Novel Pan-RAS Inhibitor with a Unique Mechanism of Action Blocks Tumor Growth in Mouse Models of GI Cancer

**DOI:** 10.1101/2023.05.17.541233

**Authors:** Jeremy B. Foote, Tyler E. Mattox, Adam B. Keeton, Xi Chen, Forrest Smith, Kristy L. Berry, Thomas Holmes, Junwei Wang, Chung-Hui Huang, Antonio B. Ward, Amit K. Mitra, Veronica Ramirez-Alcantara, Cherlene Hardy, Karrianne G. Fleten, Kjersti Flatmark, Karina J. Yoon, Sujith Sarvesh, Ganji Purnachandra Nagaraju, Dhana Sekhar Reddy Bandi, Yulia Y. Maxuitenko, Jacob Valiyaveettil, Julienne L. Carstens, Donald J. Buchsbaum, Jennifer Yang, Gang Zhou, Elmar Nurmemmedov, Ivan Babic, Vadim Gaponenko, Hazem Abdelkarim, Michael R. Boyd, Gregory S. Gorman, Upender Manne, Sejong Bae, Bassel F. El-Rayes, Gary A. Piazza

**Affiliations:** Department of Microbiology, University of Alabama at Birmingham, Birmingham AL; University of South Alabama, Mobile, AL; Department of Drug Discovery and Development, Harrison College of Pharmacy, Auburn University, Auburn, AL; Department of Gastroenterological Surgery, Oslo University Hospital, The Radium Hospital, Oslo, Norway and Institute of Clinical Medicine, Faculty of Medicine, University of Oslo, Oslo, Norway; Department of Pharmacology and Toxicology, University of Alabama at Birmingham, Birmingham, AL; Department of Hematology and Oncology, University of Alabama at Birmingham, Birmingham, AL; Department of Obstetrics and Gynecology, University of Alabama at Birmingham, Birmingham, AL; University of Connecticut, Mansfield, CT; Georgia Cancer Center, University of Augusta, Augusta, GA; Celaris Bio LLC, San Diego, CA; Department of Biochemistry and Molecular Genetics, University of Illinois, Chicago, IL; ADT Pharmaceuticals LLC, Orange Beach, AL; Department of Pharmaceutical, Social and Administrative Sciences, McWhorter School of Pharmacy, Samford University; Birmingham, AL; Department of Pathology, University of Alabama at Birmingham, Birmingham, AL; Division of Preventive Medicine, University of Alabama at Birmingham, Birmingham, AL

**Keywords:** KRAS, pan-RAS inhibitor, pancreatic cancer, colorectal cancer, tumor immune microenvironment

## Abstract

Here, we describe a novel pan-RAS inhibitor, ADT-007, that potently inhibited the growth of RAS mutant cancer cells irrespective of the RAS mutation or isozyme. RAS^WT^ cancer cells with GTP-activated RAS from upstream mutations were equally sensitive. Conversely, RAS^WT^ cancer cells harboring downstream BRAF mutations and normal cells were essentially insensitive to ADT-007. Sensitivity of cancer cells to ADT-007 required activated RAS and dependence on RAS for proliferation, while insensitivity was attributed to metabolic deactivation by UDP-glucuronosyltransferases expressed in RAS^WT^ and normal cells but repressed in RAS mutant cancer cells. ADT-007 binds nucleotide-free RAS to block GTP activation of effector interactions and MAPK/AKT signaling, resulting in mitotic arrest and apoptosis. ADT-007 displayed unique advantages over mutant-specific KRAS and pan-KRAS inhibitors, as well as other pan-RAS inhibitors that could impact *in vivo* antitumor efficacy by escaping compensatory mechanisms leading to resistance. Local administration of ADT-007 showed robust antitumor activity in syngeneic immune-competent and xenogeneic immune-deficient mouse models of colorectal and pancreatic cancer. The antitumor activity of ADT-007 was associated with the suppression of MAPK signaling and activation of innate and adaptive immunity in the tumor immune microenvironment. Oral administration of ADT-007 prodrug also inhibited tumor growth, supporting further development of this novel class of pan-RAS inhibitors for RAS-driven cancers.

**SIGNIFICANCE:** ADT-007 has unique pharmacological properties with distinct advantages over other RAS inhibitors by circumventing resistance and activating antitumor immunity. ADT-007 prodrugs and analogs with oral bioavailability warrant further development for RAS-driven cancers.

## INTRODUCTION

Colorectal cancer (CRC) and pancreatic ductal adenocarcinoma (PDA) are projected to cause 53,010 and 51,750 deaths in the US during 2024, respectively [1]. The 5-year survival rates for CRC and PDA are estimated to be 64% and 13%, respectively [1]. Over 50% of CRC and 90% of PDA patients harbor gain-in-function mutations in KRAS genes that are associated with poor prognosis, making the development of novel RAS inhibitors an urgent unmet medical need [2].

For many cancers, RAS mutations initiate tumorigenesis in cooperation with the acquisition of mutations in other oncogenes or inactivating mutations in tumor suppressor genes such as p53 [3], SMAD4 [4], APC [5], or p16 [6], driving the progression from precancerous lesions to malignant disease. Signaling through mutant RAS or activated wild-type RAS (RAS^WT^) is critical for transformation [7], cancer cell proliferation and survival [7, 8], chemoresistance [9], and metastasis [5, 10]. RAS mutations also promote immunosuppression within the tumor immune microenvironment (TiME) through various mechanisms, including increased acute inflammatory cytokines (IL-6, IL-8) [11], priming of immunosuppressive T-regs [12], MDSCs [13], downregulation of MHCI [14], increased expression of programmed cell death ligand 1 (PD-L1) [15], and suppression of intra-tumoral interferon (IFN) signaling [16]. Cumulatively, oncogenic RAS-mediated reprograming of the TiME results in weakened recognition and reduced cytotoxic T and NK cell functions supporting tumor progression.

The recent development of mutant-specific KRAS inhibitors directed against cancers harboring KRAS mutations at the G12C and G12D codons renewed optimism for selective targeting of KRAS-driven malignancies [17]. Clinical trials evaluating sotorasib (AMG-510) and adagrasib (MRTX-849) selective for the KRAS^G12C^ mutation have reported efficacy for NSCLC patients with mutant KRAS^G12C^ but a lower response rate for patients diagnosed with KRAS^G12C^ mutant CRC [18], possibly resulting from treatment related adaptive resistance [19] involving new activating mutations in RAS isozymes (K/N/HRAS) or activation of RAS^WT^ isozymes from upstream receptor tyrosine kinases (RTK) [20]. In addition, the presence of different RAS mutations in the same tumor and the role of RAS^WT^ proteins in oncogenesis [7] expands the potential efficacy advantages of RAS inhibitors capable of inhibiting both mutant RAS and hyper-activated RAS^WT^.

Here, we characterize a novel pan-RAS inhibitor, ADT-007, with potential advantages over other RAS inhibitors to circumvent resistance and activate antitumor immunity. ADT-007 emerged from screening a compound library that shares the indene scaffold of the nonsteroidal anti-inflammatory drug (NSAID), sulindac, previously reported to selectively inhibit the formation of RAS mutant tumors in a rat model of chemical-induced mammary tumorigenesis [21] and directly bind RAS to inhibit MAPK signaling, albeit with low potency and confounding cyclooxygenase inhibitory activity [22, 23]. We determined the anticancer activity of ADT-007 using cancer cell lines representative of the complex mutational landscape of RAS-driven cancers and immune modulatory potential with mouse tumor models having different immune compositions representative of the heterogeneity in patients with RAS-driven cancers.

## MATERIAL AND METHODS

### Chemical synthesis

ADT-007, (*Z*)-2-(5-fluoro-1-(4-hydroxy-3,5-dimethoxybenzylidene)-2-methyl-1H-inden-3-yl)-N-(furan-2-ylmethyl)acetamide, and ADT-316, (*Z*)-4-((5-fluoro-3-(2-((furan-2-ylmethyl)amino)-2-oxoethyl)-2-methyl-1H-inden-1-ylidene)methyl)-2,6-dimethoxyphenyl 4-methylpiperazine-1-carboxylate hydrochloride, were synthesized as follows: To a 250-mL round-bottom flask equipped with a gas outlet, (Z)-2-(1-(4-acetoxy-3,5-dimethoxybenzylidene)-5-fluoro-2-methyl-1H-inden-3-yl) acetic acid (10.3 g, 25.00 mMol) was added to 180 mL of anhydrous dichloromethane, followed by carbonyl diimidazole (CDI) (4.25 g, 26.25 mMol). The mixture was stirred for 30 min followed by adding furfuryl amine (2.68g, 26.25 mMol) and anhydrous pyridine (20 mL). The solution was stirred at room temperature for 48 h and washed with 0.2N HCl (120 mL) and water (120 mL x 2). The organic layer was concentrated, and the residue dissolved in 300 mL of methanol, followed by adding ammonium in methanol (7N, 100 mL). The red mixture was stirred vigorously overnight and filtrated. The solid was dissolved in 400 mL of ethyl acetate, treated with 1N HCl, washed with brine (150 mL x 3), and dried over anhydrous sodium sulfate. After filtration, the solution was concentrated and recrystallized from acetone/ethyl acetate to make ADT-007 as a yellow crystal. The mother liquor was concentrated and purified by silica gel column chromatography, followed by the above recrystallization procedure to yield the second batch (9.54 g combined, 85% yield). Treatment of ADT-007 with 4-methylpiperazine-1-carbonyl chloride in anhydrous pyridine at 50°C overnight, followed by chromatography purification and acidification, afforded ADT-316 HCl as a yellow crystal.

### Mice

Eight-week-old female C57BL/6 mice (000664), male and female NOD.Cg-*Prkdc^scid^ Il2rg^tm1Wjl^*/SzJ (NSG, 00557) and female B6.129S7-*Rag1^tm1Mom^*/J (Rag 1^-/-^, 02216) mice were purchased from the Jackson Laboratory. Eight-week-old female Balb/c mice were purchased from Charles River Laboratories (028). Female B6.Ifng/Thy1.1 KI (IFN-γ.Thy1.1) and B6.Cg-Tg-IL2^tm1(eGFP)/Weav^ (IL-2.GFP) knock-in cytokine reporter mice were kindly provided by Dr. Casey Weaver (University of Alabama at Birmingham, Birmingham, AL) [24]. Six-eight-week-old female athymic foxn1 nude mice were bred at the Department of Comparative Medicine, Oslo University Hospital (OUH).

### Cell lines and patient-derived xenografts

Murine PDA cell lines were derived from tumor-bearing Kras^LSL-G12D/+^; Trp53^LSL-R172H/+^; Pdx1-Cre; Rosa26^YFP/YFP^ (7160c2, 2838c3) and Kras^LSL-G12D/+^; Trp53^L/+^; Pdx1-Cre (5363) mice and cultured in DMEM complete media as previously indicated [25]. Human tumor cell lines (BxPC-3, AsPC-1, CFPAC-1, MIA PaCa-2, PANC-1, SW-1990, HT-29, HCT-116, A549, SW480, MDA-MB-231, Hs 578T, SK-MEL-2, NCI-H1299, OV-90, H-1975, U-87 MG, SK-OV-3), and the murine tumor lines (CT26 colon, B16 melanoma, and Lewis lung cancer) were purchased from ATCC and maintained according to ATCC guidelines. Human OVCAR-8 cell line was purchased from DCTD Tumor Repository (National Cancer Institute at Frederick) and maintained according to Tumor Repository guidelines. MKN-1 cells were purchased from Accegen Biotech (Paoli, PA). Normal human airway epithelial cells (NHAEC) were purchased from Lonza Bioscience and cultured using bronchial epithelial growth medium per the supplier’s protocol. Normal human colon mucosal (NCM460) cells were obtained from InCell Corporation and maintained in medium/nutrient mixture with 10% fetal bovine serum (FBS) for less than 12 passages, according to the supplier’s protocol. The human gall bladder adenocarcinoma expressing KRAS^G12V^ was derived from an 80-year-old male patient with a history of pancreatic cancer. Diagnosed initially as PDA, the tumor was reclassified as a gall bladder adenocarcinoma. This patient-derived xenograft (PDX) model was serially passaged in NSG mice. The CRC PDX models were established by implanting peritoneal metastasis samples collected at the time of cytoreductive surgery – hyperthermic intraperitoneal chemotherapy (CRS-HIPEC) into the peritoneal cavity of nude mice. The PMP-1 model was established from a patient with an appendiceal primary lesion, while PMCA-1 was from a patient with a primary mucinous rectal carcinoma [26, 27]. The PC244 model was established from a peritoneal metastasis specimen from a 69-year-old female with a poorly differentiated mucinous adenocarcinoma of the cecum. Targeted next-generation sequencing of PC244 was performed as previously described [28].

### Tumor cell transfections

BxPC-3 and HT-29 cells were initially exposed to lentiviral particles containing a mutant HRAS^G12V^ or *KRAS*^G12C^ construct for 48 h in a 6-well plate at 37°C in a humidified incubator with 5% carbon dioxide. The media was then removed and replenished with fresh media containing HA-*HRAS*^G12V^ or HA-*KRAS*^G12C^ lentiviral particles. After another 48 h, the media was removed, and fresh media containing 5 µg/mL puromycin was added to the well. Surviving cells were expanded in a 10 cm dish, and expression of the construct was confirmed by western blotting (WB) with an HA tag antibody (Cell Signaling Technology).

HEK 293FT (Invitrogen, # R70007) cell line was transfected with either pBabe-puro-HA-KRAS^G12C^ (Addgene, #58901) or pBabe-puro-Empty (Addgene, #1764) retroviral constructs along with pCL-10A1 retrovirus packaging vector (Novus Biologicals, # NBP2-29542). Following a 48-h incubation at 37°C, culture supernatants containing viral particles were collected and mixed with PEG-itTM (SBI, #LV810A-1) virus precipitation reagent (5X) followed by overnight incubation at 4°C. Supernatants were centrifuged at 1500 x g for 30 min, and white pellets of retroviral particles containing either pBabe-puro-HA-KRAS^G12C^ or pBabe-puro-Empty retroviral construct were collected and resuspended in sterile phosphate-buffered saline (PBS). BxPC-3 cells were transduced with retroviral particles in a 6-well plate using polybrene transfection reagent (EMD Millipore, TR-1003-G) and incubated for 48 h at 37°C. The media was replaced with fresh media containing 5 µg/mL puromycin. Surviving cells were selected, expanded, and confirmed for KRAS^G12C^ expression by WB using an HA tag antibody (Cell Signaling Technology).

### *In vitro* drug treatments and EGF stimulation

For *in vitro* ADT-007 studies, cell cultures were incubated with increasing concentrations (0.1, 0.5, 1, 5, 10, 50, 100, 250, 500, 1000, 10000 nM) of ADT-007, AMG-510 (Selleck Chemicals), BI-2865 or RMC-6236 (MedChemExpress), MRTX849 (Selleck Chemicals) or vehicle (DMSO) for 24 – 72 h depending on the assay. In some experiments, recombinant mouse EGF (Gibco) was added 10 min before harvesting protein lysates from treated tumor cell lines to assess the impact of upstream receptor tyrosine kinase signaling on RAS-MAPK and RAS-PI3K pathways.

### Cell growth assays

To determine 2D growth IC_50_ values, 5 x 10^3^ tumor cells were plated in triplicate using 96-well flat-bottom plates. To determine 3D growth IC_50_ values, 1 x 10^3^ tumor cells were plated in triplicate using 96-well round bottom non-binding plates. Cultures were incubated with various concentrations of ADT-007 (0.1. 0.5. 1, 5, 10, 50, 100, 250, 500, 1000, 10000 nM) and cultured for an appropriate period depending on cellular growth kinetics as indicated in figure legends. At 72-96 h (2D) or 8-10 days (3D spheroid), cells were processed according to the manufacturer’s recommendations to assess intra-cellular ATP levels as a measure of cellular proliferation and viability using CellTiter-Glo ATP content assay (CTG assay) (Promega). Nonlinear dose-response curves showing the effects of compound concentration on colony count were generated using GraphPad Prism (GraphPad Software).

### Mouse liver microsome stability assays

ADT-007 (1 µM) was incubated with mouse liver microsomes along with a NADPH regeneration solution and UGT cofactors (glucuronidation) in a phosphate buffer (pH 7.4) at 37°C in a shaking water bath. Samples were taken at 0, 5, 15, 20, 60 and 120 min, quenched with acetonitrile and analyzed by LC-tandem mass spectrometry to measure the remaining amount of ADT-007 as a function of time. Peak area ratios of ADT-007 to an internal standard (terfenadine) were calculated at each time point and data were expressed as percent of time zero. Samples at each time point were also analyzed in a Q1 and triggered independent data acquisition scan to determine the identity of the metabolites. The data were processed using LightSight metabolite ID software. The primary metabolite was found to be a mono-glucuronide conjugate of ADT-007.

### *In vitro* intracellular compound analysis

Triplicate samples of 60% confluent HT-29 and HCT-116 cells were treated with 100 nM ADT-007 for 16h in 15 cm plates. Treatment medium was collected and snap frozen. Cell pellets were collected by scraping and weighed in tared tubes. Concentration of ADT-007 and relative ratio of ADT-007 glucuronide (m/z 446+193) were quantified by LC-MS/MS as described above.

### Gene expression database analysis

Transcriptomic (RNAseq data) for most of the cell lines in Table 1 is publicly available in Cancer Cell Line Encyclopedia accessed via the DepMap Portal [29]. The Student’s t-test was used to identify significant differences in 19,221 TPM (transcripts per million) +1 normalized transcripts between the two categories of cell lines. Due to the modest number of resistant lines available, a threshold of p < 0.01 was chosen, which identified 719 transcripts and were further reduced to 254 unique transcripts which expressed at least a 5-fold difference.

**Table 1.** ADT-007 sensitivity is associated with high activated RAS and low UGT superfamily enzyme levels. Growth inhibitory (IC_50_) values following 72 h of treatment (CTG assay). TPM normalized expression of the most significant differentially expressed UGT1A10 enzyme from published data (Cancer Cell Line Encyclopedia; DepMap Portal; Broad Institute). * Expression data not available, † TPM normalized UGT2B7 expression in SK-MEL-28.

| Type | Tissue Origin | Cell Line | Mutation Status | IC<sub>50</sub> (nM) | UGT1A10 |
| --- | --- | --- | --- | --- | --- |
| Mutant RAS, sensitive | Pancreatic | MiaPaCa2 | KRAS G12C | 2.1 | 0.263 |
|  | Pancreatic | Panc1 | KRAS G12D | 2.4 | 0.043 |
|  | Pancreatic | SW1990 | KRAS G12D | 6.3 | 0.138 |
|  | Lung | A549 | KRAS G12S | 6.8 | 0.070 |
|  | Pancreatic | CFPAC1 | KRAS G12V | 5.8 | 2.63 |
|  | Colon | SW480 | KRAS G12V | 6.5 | 0.58 |
|  | Breast | MDA231 | KRAS G13D | 3.6 | 0.043 |
|  | Colon | HCT116 | KRAS G13D | 4.7 | 0 |
|  | Ovarian | OVCAR8 | KRAS P121H | 5.8 | 0.057 |
|  | Breast | HS578 | HRAS G12D | 3.9 | 0.098 |
|  | Skin | SKMEL2 | NRAS Q61K | 2.1 | 0 |
|  | Lung | H1299 | NRAS Q61K | 2.4 | 0 |
| WT RAS, resistant | Normal Lung | NHAEC | WT RAS | 19100 | * |
|  | Normal Colon | NCM460 | WT RAS | 50000 | * |
|  | Gastric | MKN-1 | Amplified WT KRAS | 4.4 | 0.49 |
| Downstream mutation (WT RAS) resistant | Ovarian | OV90 | WT RAS/mut BRAF | 350 | 4.57 |
|  | Colon | HT29 | WT RAS/mut BRAF | 512 | 6.97 |
|  | Pancreatic | BXPC3 | WT RAS/mut BRAF | 2500 | 7.75 |
| Upstream RAS activation sensitive | Lung | H1975 | WT RAS/mut EGFR | 3.9 | * |
|  | Skin (mouse) | B16 | WT RAS/mut PDGFR | 5.8 | * |
|  | Brain | U87MG | WT RAS/mut NF1 | 1.6 | 0 |
|  | Ovarian | SKOV3 | WT RAS/mut NF1 | 6.7 | 0.202 |
| Amplified WT RAS, sensitive | Gastric | MKN-1 | Amplified WT KRAS | 4.4 | 0.490 |

### NMR Spectroscopy

Recombinant KRAS^WT^ was dialyzed 2x for 24 h (12 h each in 1L 50 mM Tris-Citrate pH=6.5, 50 mM NaCl, 10 mM β-mercaptoethanol) at 4°C following treatment with 10 mM EDTA to remove Mg^2+^ and the bound nucleotide. To record ^1^H- ^15^N heteronuclear single quantum coherence (HSQC) NMR spectra of nucleotide-free KRAS (nf-KRAS), 900 MHz and 600 MHz Avance spectrometers (Bruker) equipped with cryogenic probes were used. Each experiment was performed at 25°C in 50 mM Tris-Citrate pH=6.5, 50 mM NaCl, 10 mM β-mercaptoethanol containing 10% ^2^H_2_O. The chemical shift assignments were taken from the Biological Magnetic Resonance Data Bank (BMRB) database (http://www.bmrb.wisc.edu) using the BMRB accession numbers 18529, 25114, and 26635. Chemical shift perturbations higher than the average sum and one standard deviation (SD) were considered statistically significant.

### Nucleotide Exchange Assays

The fluorescent GTP analog MANT-GTP was purchased from Millipore Sigma. For each reaction, 5 μL of 5 μM purified GDP-bound KRAS in reaction buffer (40 mM Tris-HCl, 100 mM NaCl, 20 mM MgCl_2_, 1 mg/mL BSA, pH 7.5) and 2.5 μL of 200 mM EDTA were added to a microcentrifuge tube to make a master mix. In some reactions, water was used in place of EDTA to control the release of GDP. 7.5 μL of master mix per well was added to a 384-well plate for each reaction. Compound (5 μL) was added to each respective well at five times the desired concentration or vehicle in the reaction buffer. After 2.5 min for equilibration at 26°C in a Bio-Tek Synergy H4 microplate reader, 12.5 μL of 1.5 μM MANT-GTP in reaction buffer was added to each well for a final concentration of 750 nM MANT-GTP, 1 μM RAS, 20 mM EDTA, and the desired concentration of compound or vehicle. In the case of compound preincubation, the plate was incubated for 1 h at 26°C before MANT-GTP addition. Fluorescence was measured at an excitation wavelength of 360 nm and an emission wavelength of 460 nm in a Bio-Tek Synergy H4 microplate reader. Kinetic curves were generated with GraphPad Prism.

### Cellular thermal stability assays

Experiments to determine ADT-007 binding to RAS in cells were performed by Cellaris Bio LLC using HCT-116 and HT-29 cells cultured in RPMI-1640 medium supplemented with 10% FBS. To determine the melting profile of KRAS, fresh cell lysates were prepared in non-denaturing buffer and cleared *via* centrifugation at 5,000 rpm for 15 min. Cleared lysates were dispensed into 96-well PCR (equivalent of approx. 6,000 cells/well/50 µl), then subjected to the temperature gradient (38-65°C) for 10 min. Subsequently, centrifugation was performed at 14,000 rpm to sediment the unstable protein content. Following WB of the soluble fraction, bands of stable KRAS were quantified, normalized to DMSO, and plotted using GraphPad Prism. Nonlinear regression analysis was used to fit a sigmoidal dose-response curve with variable slope to generate EC_50_ values for ADT-007 binding to KRAS.

Additional cellular thermal stability experiments used a Micro-Tag enzyme complementation method. A KRAS^G12C^ Micro-Tag construct was engineered with a 15-amino acid tag at the N-terminus. The construct was transiently expressed in HEK293 human embryonic kidney cells. Two days post-transfection, cells were suspended, washed with Tris Buffered Saline (TBS: 20mM Tris HCl pH 7.4, 150mM NaCl), and resuspended at 1000 cells/µl in TBS and 50,000 cells aliquoted to wells of a PCR plate. Increasing concentrations of ADT-007 were added and incubated with cells for 45 min followed by heating for 10 min at 55°C and a 1 min cool-down on ice. An equal volume of 0.5% Triton-X-100 in TBS was added, and cells lysed for 10 min on ice. Fluorescence was measured and plotted with the concentration of ADT-007 on a semi-log scale using GraphPad Prism 9.0 software. Nonlinear regression analysis was used to fit a sigmoidal dose-response curve with variable slope to generate EC_50_ values for ADT-007 binding to KRAS.

### RAS activation assays (RAS-RBD pulldown assays)

Cells were grown to 80 percent confluence using complete growth medium, then washed with 1 mL of PBS and lysed on ice for 5 min (25 mM Tris HCl, 150 mM NaCl, 5 mM MgCl_2_, 1% NP-40, 5% glycerol, pH 7.2) with Halt Protease and Phosphatase Inhibitor Cocktail (Thermo Fisher Scientific), then clarified at 16,000 RCF 15 min at 4°C. Protein concentration of the supernatant was normalized using the DC protein assay (Bio-Rad). 200 µg of whole cell lysate was incubated with GST-RAF-RBD (Addgene) bound to glutathione agarose beads at 4°C for 1 h. Bound active RAS was collected by centrifugation at 6,000 RCF for 30 seconds at 4°C, washed thrice with lysis buffer, then eluted with SDS sample buffer and subsequently detected by WB with a pan-RAS antibody or isozyme-specific antibody as described below.

### Western blotting

Tumor or cell lysates were generated after treatment with 500 µL RIPA lysis buffer (Thermo Scientific, 89900) with protease and phosphatase inhibitors (Thermo Scientific, 78440), boiled for 5 min, and equal protein concentrations were loaded, and run on a 12% SDS-PAGE. Gels were transferred to a PVDF membrane, which was blocked with either 5% non-fat dry milk in TBS-T (TBS with 0.1% Tween 20) or 5% BSA in TBS-T, depending on the blocking agent used for the primary antibodies, for 1 h at room temperature. Membranes were incubated overnight at 4°C on a rocker with primary antibodies **(Supplemental Table 1)** according to the manufacturer’s instructions and then developed with HRP-conjugated secondary antibodies (Cell Signaling Technology) for 1 h at room temperature on an orbital shaker. Membranes were developed using SuperSignal West Enhanced Chemiluminescent substrate (Thermo Fisher Scientific) and imaged using a ChemiDoc Imaging System (Bio-Rad).

### *In vivo* efficacy experiments

#### Murine tumor models

A single cell suspension of murine CT26 CRC or 2838c3, 7160c2, and 5363 PDA KRAS^G12D^ cells was prepared in PBS and kept on ice until injection. Syngeneic mouse CRC and PDA tumor cells (1×10^6^/mouse) were injected subcutaneously (SQ) into 6-8-week-old Balb/c or C57BL/6 mice, respectively. When tumors reached an average tumor volume of 150 mm^3^, mice began receiving daily intra-tumoral (10 mg/kg, QD, 7 days/week for 13 days) or peri-tumoral (5 mg/kg, BID, 7 days/week for 13-24 days) treatment with ADT-007. In all experiments tumor growth was assessed by caliper measurements (2-3/week), and tumor volumes (TV = W x L^2^/2) were calculated throughout the study. No overt signs of toxicity (body weight loss or organ [lungs, heart, liver, GI tract] pathology) were noted with local administration.

#### PDX CRC tumor models

mucinous PMCA-1, PMP1, and PC244 PDX colorectal adenocarcinomas were implanted intraperitoneally (IP) in six-eight-week-old female athymic foxn1 nude mice. Treatment with ADT-007 (IP, 0.33, 0.67, 1.25 and/or 2.5 mg/kg, BID, 5 days/week for 18-23 total days of treatment) was initiated 24 h after implantation (n = 5-6 mice/group) and mice were followed by MRI for 24 to 50 days after implantation (6-31 days post-treatment cessation).

#### PDX GI tumor model

KRAS^G12V^ PDX gall bladder adenocarcinoma was implanted SQ in eight-week-old male NSG mice. When tumors reached an average tumor volume of 150 mm^3^, mice began receiving daily peri-tumoral (5 mg/kg, BID, 7 days/week for 13 days, n = 5 mice/group) treatment with ADT-007. Tumor growth was assessed by caliper measurements (2-3/week), and tumor volumes were calculated throughout the study.

All experiments were randomized but not blinded. For animal studies performed at the University of Alabama at Birmingham (UAB) and the University of South Alabama all procedures were performed under approved protocols according to the Institutional Animal Care & Use Committee (IACUC) guidelines and Animal Resources Program of UAB. All cell lines and PDX models were tested against an extended panel of excluded pathogens, including mycoplasma, before implantation into mice. For animal studies performed at OUH Institute for Cancer Research (Norwegian Radium Hospital, Oslo, Norway) all procedures and experiments involving animals were approved by the Norwegian Food Safety Authority (application ID #18209) and were conducted according to the recommendations of the European Laboratory Animals Science Association and the ARRIVE guidelines.

### Flow Cytometry

Mouse tumors were digested in a 1 mg/mL collagenase IV and 0.1 mg/mL DNAase 1 (Worthington Biochemical) in HBSS for 45 min at 37 °C with intermittent shaking. Samples were washed with media and filtered through a 70 μM strainer generating suspensions of single cells. Cells were labeled with primary fluorophore-conjugated antibodies and a live/dead stain (**Supplemental Table 2**) for 30-60 min at 4°C, washed, and resuspended in flow buffer for downstream immune cell analyses (**Supplemental Figures 9A and 10A**). Cytokine expression from single-cell suspensions of spleen and tumor was quantified as follows: cells were plated at a concentration of 10^6^ cells in 1 mL of RP-10 media and stimulated for 5 h at 37°C using a 1x solution of a Cell Activation Cocktail containing PMA ionomycin, and Brefeldin A (BioLegend). Conversely, cells were cultured for 5 h at 37°C using a solution containing 5 µg of Brefeldin A (BioLegend) as an unstimulated control. After cell surface staining, cells were fixed in 4% paraformaldehyde for 45 min at 4°C then washed with 1x Perm/Wash (BD, 554723). For intracellular staining in tumor and spleen samples, cells were stained with an antibody cocktail-specific intracellular cytokines (IL-2, IFNγ, TNFα, and granzyme B) in Perm/Wash solution for 60 min at 4°C. Data were acquired on a Symphony A5 flow cytometer, with analysis performed using FlowJo version 10.7.2.

### Statistical Analyses

GraphPad Prism (version 9.1.2) was used for statistical analyses and graphical representation with data presented as either means ± standard deviation (SD) or standard error of the mean (SEM). Two-tailed Student’s or Welch’s t-test and two-way Analysis of Variance (ANOVA) with multiple corrections (Tukey’s or Sadik’s method) were performed to determine the statistical significance between groups. To calculate IC_50_ values, compound concentrations were log-transformed, and data normalized to controls to generate a *log*(inhibitor) *vs*. response (four-parameter nonlinear dose-response). A p < 0.05 was considered statistically significant.

### Data Availability

Data generated in this study will be made available upon request from the corresponding author.

## RESULTS

### ADT-007 potently and selectively inhibits the growth of cancer cells with mutated or activated RAS

An indene library was synthesized and screened for differential growth inhibitory activity by comparing sensitivity of CRC HCT-116 cells harboring a KRAS^G13D^ mutation *vs*. HT-29 cells with RAS^WT^ but mutant BRAF^V600E^. A lead compound, ADT-007 (**Figure 1A**), and several analogs emerged following chemical optimization for potency and selectivity. ADT-007 inhibited HCT-116 growth with a 5 nM IC_50_, while HT-29 cells were ∼100-fold less sensitive (**Figure 1B**). RAS-RBD pulldown assays confirmed high activated RAS (RAS-GTP) levels in HCT-116 cells, while RAS-GTP was undetectable in HT-29 cells. Despite differences in RAS-GTP levels, HCT-116 and HT-29 cells depended on MAPK signaling for growth, as both were sensitive RAF, MEK, and ERK inhibitors (not shown). The requirement of mutant RAS for sensitivity to ADT-007 was determined by transfecting HT-29 cells with HRAS^G12V^. ADT-007 treatment of RAS-transfected HT-29 cells resulted in a 24 nM IC_50_ in growth assays compared with 549 and 730 nM IC_50_ values for parental and vector-control cells, respectively (**Figure 1C**).

**Figure 1.**
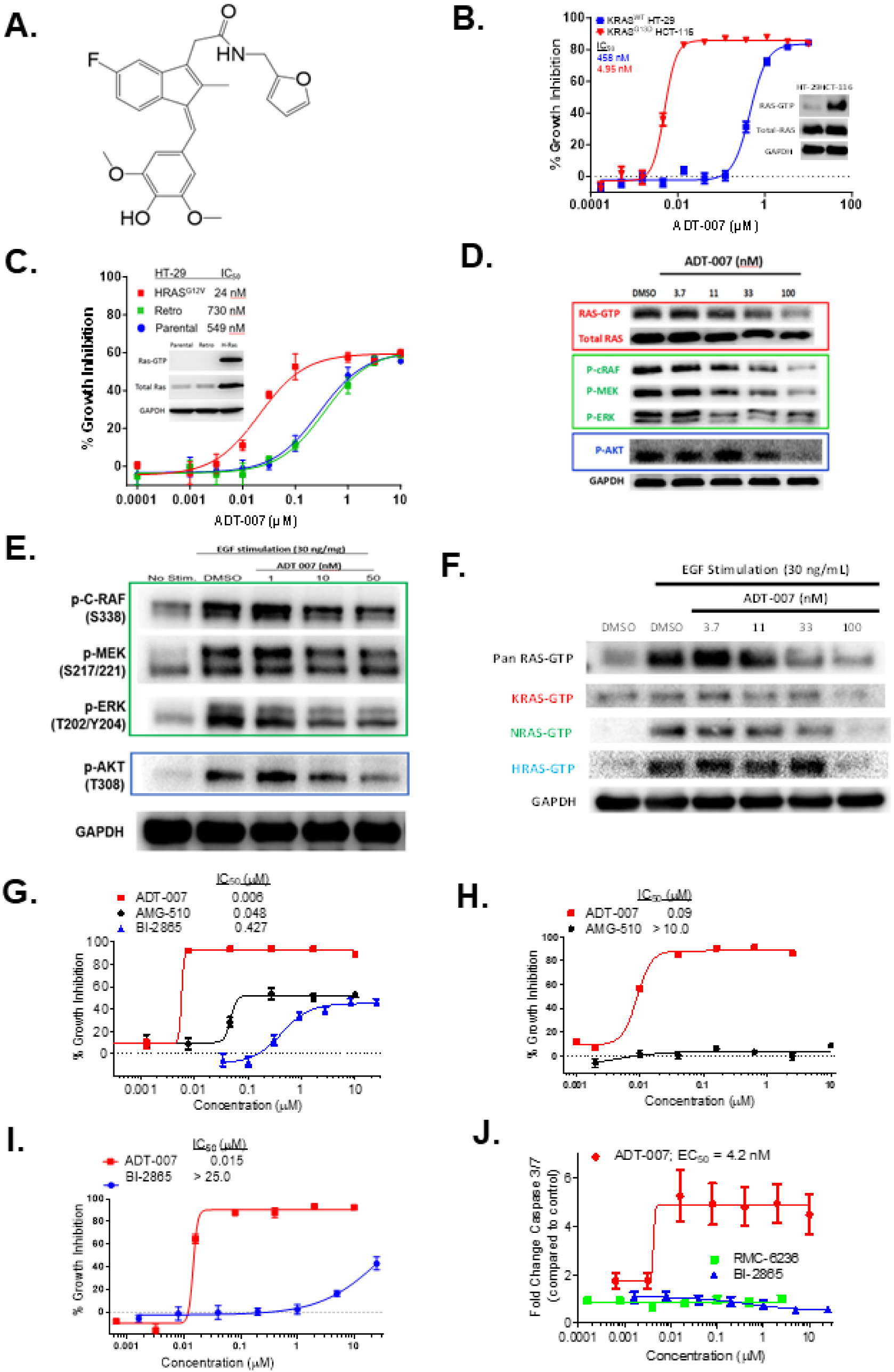
ADT-007 selectively inhibits growth of RAS mutant cancer cells by blocking activated RAS and MAPK/AKT signaling and superior in comparison with other RAS inhibitors. **(A)** Chemical structure of ADT-007. **(B)** ADT-007 inhibition of KRAS^G13D^ HCT-116 vs. KRAS^WT^, BRAF^V600E^ HT-29 CRC cell growth following 72 h of treatment (CTG assay). Quadruplicate samples ± SEM. Inset: Activated RAS-GTP-RBD pulldown evaluated by WB using pan-RAS antibodies. Total RAS and GAPDH served as loading controls. **(C)** Ectopic expression of HRAS^G12V^ sensitized HT-29 cells to ADT-007 growth inhibition (quadruplicates ± SEM). Inset: Activated RAS in untreated parental, vector control (retro), and transfected cells (H-Ras). (**D**) HCT-116 KRAS^G13D^ CRC cells were treated with increasing concentrations of ADT-007 overnight in complete medium (10% FBS). Activated RAS-GTP pulldowns, phospho(P)-cRAF, P-MEK, P-ERK, P-AKT, and total RAS and GAPDH as loading controls were detected by WB. Data represent 3 independent experiments. (**E, F**) Serum-starved MIA PaCa-2 KRAS*^G12C^* PDA cells were treated overnight with vehicle or increasing concentrations of ADT-007, then EGF stimulated (30 ng/mL, 10 min). (**E)** P-cRAF, P-MEK, P-ERK, P-AKT and GAPDH were detected by WB. Data represent two independent experiments. (**F**) Activated RAS-GTP was isolated by RAS-RBD pulldown followed by WB with pan-RAS, KRAS, NRAS, or HRAS specific antibodies and GAPDH. Data represent at least two independent experiments. (**G**) Growth inhibition of MIA PaCa-2 cells by ADT-007 *vs.* BI-2865 or AMG-510 as determined by CTG assay following 72-h treatment (**H, I**) Growth inhibition of LLC cells (KRAS^G12C^ / NRAS^Q61H^) by ADT-007 *vs.* BI-2865 or AMG-510 following 72-h treatment. (**J**) Apoptosis induction of MIA PaCa-2 cells following 48-h treatment with ADT-007, BI-2865 or RMC-6236 as measured by cleaved caspase-3/7 using a luminescence assay (Promega). Error bars represent SEM.

A panel of histologically diverse cancer cell lines was also screened for sensitivity to ADT-007 (**Table 1**). Cancer cell lines with mutations in K/N/HRAS isozymes or upstream signaling components (e.g., EGFR, PDGFR, NF1) were uniformly sensitive to ADT-007 with low nM IC_50_ values. Like HT-29 cells, other cancer cell lines with downstream BRAF mutations were essentially insensitive to ADT-007, as were normal colonocytes and airway epithelium cells.

KRAS^G12C^ MIA PaCa-2 PDA cells were highly sensitive to ADT-007 with a 2 nM IC_50_ in growth assays while RAS^WT^ BxPC-3 PDA cells had a 2500 nM IC_50_ (**Table 1 and Supplemental Figure 1A**). RAS-RBD pulldown assays confirmed high RAS-GTP in MIA PaCa-2 and three other PDA cell lines that were comparably sensitive to ADT-007, which harbor KRAS^G12V^ and KRAS^G12D^ (**Supplemental Figure 1B**) and represent the most prevalent KRAS mutations in PDA patients [30]. By comparison, RAS-GTP levels in BxPC-3 cells were undetectable. Consistent with results from HT-29 cells, transfecting BxPC-3 cells with KRAS^G12C^ induced sensitivity to ADT-007 in growth assays (**Supplemental Figure 1C-D**).

ADT-007 treatment of MIA PaCa-2 cells blocked growth in 2D monolayer or 3D spheroid cultures with comparable low nM IC_50_ values, resulting in unusually steep dose-response curves that was a characteristic feature of ADT-007 and related analogs (**Supplemental Figure 1E-F**). ADT-007 also potently blocked colony formation of KRAS mutant PDA cell lines without impacting colony formation of RAS^WT^ BxPC-3 cells (**Supplemental Figure 1G**). In addition, ADT-007 potently and selectively inhibited colony formation of HCT-116 cells without effects on HT-29 cells (**Supplemental Figure 1H**).

### ADT-007 inhibits activated RAS and MAPK/AKT signaling

ADT-007 reduced RAS-GTP and phosphorylated MAPK/AKT components in HCT-116 cells (**Figure 1D**). ADT-007 also inhibited EGF-stimulated MAPK/AKT signaling in MIA PaCa-2 cells (**Figure 1E**). ADT-007 reduction of RAS-GTP occurred if cells were grown in serum, serum-starved, or EGF stimulated (**Supplemental Figures 2A-C**). ADT-007 also decreased RAS-GTP by treating lysates from HCT-116 cells or HRAS^G12V^ transfected HT-29 cells, suggesting a direct effect of ADT-007 on RAS protein rather than on RAS signaling (not shown). ADT-007 treatment of MIA PaCa-2 cells serum-starved and stimulated with EGF also reduced levels of GTP-HRAS^WT^ and GTP-NRAS^WT^ isozymes co-expressed with GTP-KRAS^G12C^ (**Figure 1F**). As expected, EGF did not impact GTP-KRAS^G12C^ levels but activated GTP-HRAS^WT^ and GTP-NRAS^WT^ isozymes. Nonetheless, ADT-007 reduced GTP-KRAS^G12C^ to the same extent as EGF-activated RAS^WT^ isozymes.

### ADT-007 induces mitotic arrest and apoptosis

ADT-007 induced G2/M arrest in HCT-116 and MIA PaCa-2 cells following 24 h treatment at concentrations comparable to growth IC_50_ values while concentrations at least 100x higher were required to have a similar effect in HT-29 or BxPC-3 cells (**Supplemental Figure 3A-D**). Time-course experiments revealed that ADT-007 induced apoptosis of MIA PaCa-2 cells following G2/M arrest (**Supplemental Figure 3E**). ADT-007 also induced apoptosis in HCT-116 cells but not HT-29 cells (**Supplemental Figure 3F**). Mitotic arrest was confirmed by experiments showing that ADT-007 treatment of HCT-116 cells increased phospho-histone 3B levels, which coincided with reduced cell number and occurred at concentrations within growth IC_50_ values (**Supplemental Figure 3G**).

Given previous reports that other indenes bind tubulin and induce mitotic arrest [31], the effect of ADT-007 on tubulin polymerization was determined using recombinant α-tubulin. ADT-007 exhibited a weak inhibitory effect on tubulin polymerization at concentrations ∼100-fold above growth IC_50_ values (**Supplemental Figure 3H**). Similar weak effects on tubulin polymerization were observed by treating RAS mutant cancer cells with ADT-007 (not shown). Moreover, colchicine, a known tubulin inhibitor, lacked selectivity to inhibit the growth of KRAS mutant cell lines.

### ADT-007 induces cytotoxicity; other RAS inhibitors result in cytostasis

ADT-007 was compared with the KRAS^G12C^ inhibitor, AMG-510, and the pan-KRAS inhibitor, BI-2865, in growth assays involving MIA PaCa-2 cells. Both AMG-510 and BI-2865 were significantly less potent and caused less growth inhibition than ADT-007 (**Figure 1G**). Unlike ADT-007, neither compound induced mitotic arrest as determined by phospho-histone 3B levels (**Supplemental Figure 3G**). Another key difference that may be relevant to resistance was apparent in experiments using LLC cells harboring KRAS^G12C^ and NRAS^Q61H^ mutations. LLC cells were found to be insensitive to AMG-510 and BI-2865 in growth assays but retained full sensitivity to ADT-007 (**Figure 1H-I**).

We also compared ADT-007 with another pan-RAS inhibitor, RMC-6236. While RMC-6236 displayed comparable potency as ADT-007 in growth assays involving MIA PaCa-2 cells, viable cells were observed microscopically after RMC-6236, as well as BI-2865, treatment, while only rounded condensed cells were present after ADT-007 treatment (**Supplemental Figure 4A**). Consistent with experiments described above (**Supplemental Figure 3E-F**), ADT-007 induced apoptosis of MIA PaCa-2 cells with a 4 nM EC_50_. However, neither RMC-6236 nor BI-2865 induced apoptosis (**Figure 1J**). Moreover, colony formation assays revealed that ADT-007 caused near complete inhibition of colony formation, while BI-2865 and RMC-6236 were appreciably less effective (**Supplemental Figure 4B-C**). These results show fundamental differences between ADT-007, which selectively induces apoptosis (cytotoxicity) of RAS mutant cancer cells, while the growth inhibitory activity of other RAS inhibitors appears to be limited to cytostasis.

### Chemical and biological specificity of ADT-007

The chemical specificity of ADT-007 was evaluated using a closely related chemical analog, ADT-106, containing a methoxy group in place of the hydroxyl moiety. ADT-106 lacked the selectivity of ADT-007 to inhibit the growth of MIA PaCa-2 cells compared with BxPC-3 cells (**Figure 2A-B**). ADT-106 also had reduced potency with a 150 nM IC_50_. It is also noteworthy from these experiments that the potency of ADT-007 to inhibit MIA PaCa-2 growth paralleled its potency to reduce RAS-GTP levels, while ADT-106 did not impact RAS-GTP levels. These results show the requirement of the phenolic hydroxyl moiety for both the selectivity and potency of ADT-007, as well as the association between growth inhibition by ADT-007 and its ability to suppress RAS-GTP.

**Figure 2.**
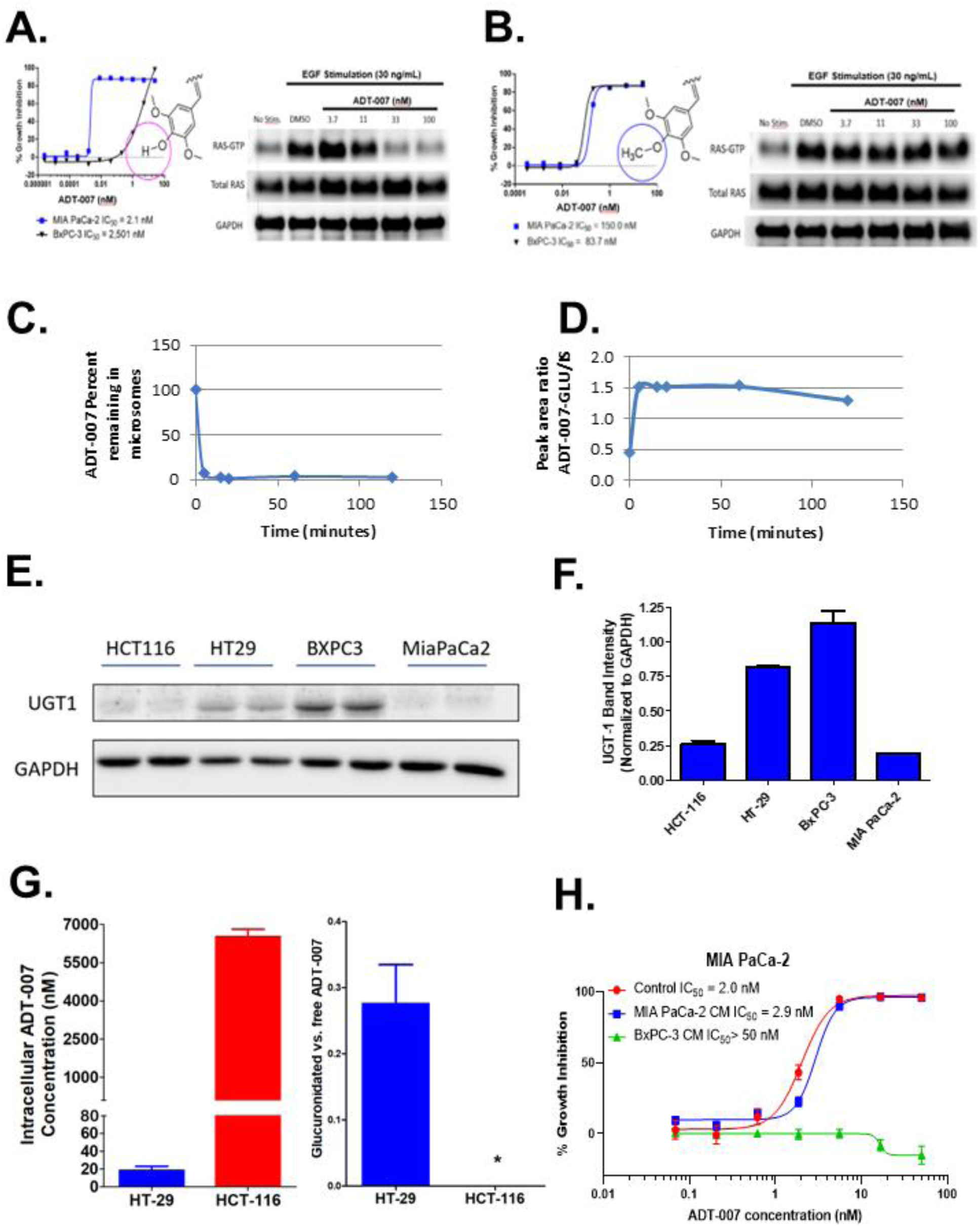
Phenolic moiety is critical for RAS selectivity of ADT-007 and renders sensitivity to metabolic deactivation. (**A**) Left panel: ADT-007 selectively inhibits the growth of KRAS^G12C^ MIA PaCa-2 vs. RAS^WT^ BxPC-3 PDA cells (72 h, CTG assay; triplicates ± SEM). (**B**) Left panel: ADT-106 lacking a phenolic OH moiety was less potent and nonselective (72 h, CTG assay; triplicates ± SEM). Right panels (**A, B**): MIA PaCa-2 cells were treated for 18 h with increasing concentrations of ADT-007 (**A**) or ADT-106 (**B**) and activated RAS-GTP-RBD pulldown evaluated by WB using pan-RAS antibodies. Total RAS and GAPDH served as loading controls. (**C**) ADT-007 is rapidly degraded by mouse liver microsomes with a corresponding increase in ADT-007-glucuronide (ADT-007-Glu) levels (**D**). (**E**) UGT1 protein expression in ADT-007 sensitive *vs*. resistant tumor cells by WB. (**F**) UGT1 protein expression and control GAPDH (WB) from (**E**) were quantified and normalized using Image J. Representative of two independent experiments. (**G**) ADT-007 and ADT-007-glucuronide levels in HCT-116 and HT-29 cell pellets following 24 h treatment with ADT-007. Levels were quantified in triplicate by LC-MS/MS (mean ± SEM), * = p < 0.05. (**H**) Conditioned medium (CM) from MIA PaCa-2 (low UGT expression) and BxPC-3 (high UGT expression) PDA cells pretreated with ADT-007 at 100 nM for 24 h was collected for growth inhibition assays involving MIA PaCa-2 cells. CM from MIA PaCa-2 cells showed similar growth inhibitory activity as the control group using freshly prepared ADT-007. CM from BxPC-3 cells resulted in complete loss of potency of ADT-007 to inhibit the growth of MIA PaCa-2 cells (96 h, CTG assay; triplicates ± SEM).

### Metabolic mechanisms impacting sensitivity to ADT-007

*Ex vivo* treatment of mouse liver microsomes with ADT-007 followed by LC-MS/MS analysis of metabolites revealed that the phenolic hydroxyl moiety is rapidly glucuronidated by UDP-glucuronosyltransferases (UGTs) for which multiple UGT isozymes exist in various cell types (**Figure 2C-D**). These results suggest that UGTs could contribute to the sensitivity of cells to ADT-007 both *in vitro* and *in vivo*. Measurement of UGT1 revealed higher protein levels in RAS^WT^ HT-29 and BxPC-3 cells compared with KRAS mutant HCT-116 and MIA PaCa-2 cells (**Figure 2E-F**). Transcriptomic data (RNAseq) analysis (Cancer Cell Line Encyclopedia) for mRNA levels of six distinct UDP-glucuronosyltransferase isozymes (UGT1A1, 1A6, 1A10, 2A1, 2B10, and 2B15) in most cancer cell lines from **Table 1** [29] provided additional evidence that RAS^WT^ cancer cell lines express higher levels of UGT isozymes compared with RAS mutant cancer cell lines (**Table 1, Supplemental Table 3**).

Consistent with these differences, measurement of ADT-007 levels by LC-MS/MS from cells treated overnight with ADT-007 resulted in 350-fold higher intracellular levels in HCT-116 cells than in HT-29 cells (**Figure 2G**). We confirmed that UGTs selectively deactivated ADT-007 in RAS^WT^ cancer cells but not in RAS mutant cells by detection of glucuronide-conjugated ADT-007 in HT-29 cells as none was detected in HCT-116 cells. In other experiments, we determined if exposure of BxPC-3 cells with high UGTs to ADT-007 could metabolize ADT-007 by collecting “conditioned medium” (CM) following 24 h treatment with ADT-007. CM from BxPC-3 cells did not inhibit MIA PaCa-2 growth, while CM from MIA PaCa-2 cells caused comparable growth inhibition as freshly prepared ADT-007 (**Figure 2H**). Similar results were obtained using CM from HT-29 cells and testing for growth inhibition of HCT-116 cells (not shown). Additional experiments showed that pretreatment with propofol, a UGT inhibitor, increased the sensitivity of BxPC-3 and HT-29 cells to ADT-007 in growth assays without significantly affecting the sensitivity of MIA PaCa-2 or HCT-116 cells (**Supplemental Table 4)**.

### ADT-007 binds RAS in cells

The ability of ADT-007 to directly bind KRAS in cells was measured using cellular thermal stability assays based on the principle that a protein maintains thermal stability when in a conformation favorably for binding to a small molecule inhibitor. Initial experiments measured the optimal temperature to denature 75% (T_agg75_) KRAS in whole cell lysates, followed by WB quantification of non-denatured KRAS levels. Treatment of HCT-116 lysates with ADT-007 increased KRAS thermal stability (T_agg75_), resulting in a 0.45 nM EC_50_ (**Figure 3A**). By comparison, ADT-007 did not affect thermal stability of KRAS in HT-29 cells. ADT-007 also did not impact thermal stability of an unrelated protein, PDE10A, in either HCT-116 or HT-29 cells known to be expressed in both cell lines [32] (not shown). Additional experiments involving ADT-007 treatment of intact HEK293 embryonic kidney cells transfected with epitope-tagged mutant KRAS^G12C^ and detection of thermal stability by WB analysis with anti-Micro-Tag antibody resulted in an 8.49 nM EC_50_ (**Figure 3B**). Consistent with experiments described above (**Figure 2A-B**) showing the importance of the phenolic hydroxyl moiety for potency and RAS selectivity, a trimethoxy ADT-007 analog did not impact thermal stability of KRAS (not shown).

**Figure 3.**
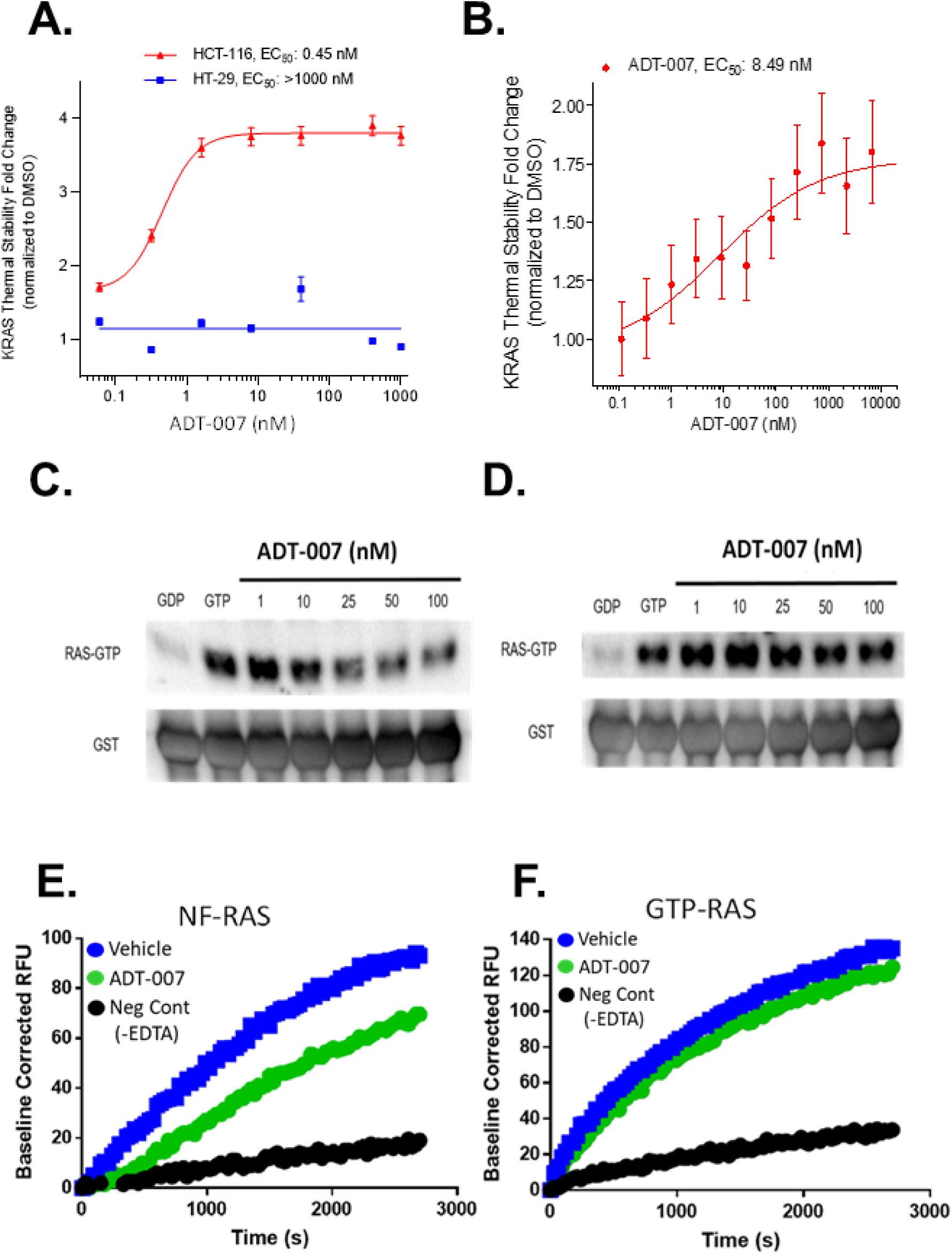
ADT-007 binds KRAS in cells and inhibits nucleotide loading and effector binding using recombinant RAS. Cellular target engagement assays show: (**A**) ADT-007 binds KRAS with high affinity in KRAS^G13D^ HCT-116 but not KRAS^WT^ HT-29 CRC cells. Graphs show levels of compound stabilized KRAS detected by WB compared with vehicle control. Curves are graphed as the average of two replicates ± SEM. (**B**) Binding affinity of ADT-007 for mutant KRAS^G12C^ was assessed by 45 min treatment of HEK293 cells expressing KRAS^G12C^-Micro-Tag construct compared with DMSO vehicle. Curve is graphed as the average of two replicates ± SEM. (**C-D**) The inhibitory activity of ADT-007 to block recombinant KRAS^WT^ activation (effector binding) in a cell-free system was evaluated by RAS-RBD pulldown assays. (**C**) ADT-007 inhibited KRAS^WT^-RAF activation before GTP addition. (**D**) ADT-007 did not significantly inhibit KRAS^WT^ activation when added after GTP loading. (**E-F**) Kinetics of GTP binding to KRAS were measured using a fluorescent GTP analog (Mant-GTP). ADT-007 reduced the rate of Mant-GTP binding to KRAS if pretreated with EDTA to chelate Mg2+ (**E**), but not if added after non-hydrolysable GTPγS before incubation with ADT-007 (**F**). Data are representative of at least two independent experiments.

### Molecular modeling studies

Molecular docking studies using human GDP-bound KRAS^WT^ (PDB ID 4OBE) [33] were conducted using molecular dynamics and induced-fit to identify potential binding sites for ADT-007 to bind RAS. In simulations of Mg^+2^ present or absent, the best GLIDE docking scores (-11.08 and -12.06, respectively) were calculated with ADT-007 binding to a site partially overlapping the nucleotide-binding domain that was associated with an altered conformation of the switch-1 region **(Supplemental Figure 5A-D)**. These results include the use of explicit water molecules that were especially critical for ADT-007 binding to 4OBE (KRAS^WT^ Mg^+2^ present). When Mg^+2^ was present, the phenolic hydroxyl moiety of ADT-007 was predicted to H-bond with Ser17 through a water bridge and with Glu31 through the amide side chain projecting into solvent space. If Mg^+2^ was absent, the phenolic hydroxyl was predicted to interact with Asp119 via a water bridge and Ala146 by a direct H-bond. Whether Mg^+2^ was present or absent, the GLIDE docking score of GDP to bind RAS (-9.6 and -8.80, respectively) was considerably less favorable than ADT-007 (not shown). These results suggest that a conformational change induced by the binding of ADT-007 may decrease the affinity of RAS to bind nucleotide since in the docking score for GDP to bind native RAS is -15.15. The results also indicate that there may be a difference in how ADT-007 binds RAS depending on whether RAS is in a Mg^+2^ bound (nucleotide-loaded) conformation or Mg^+2^ free conformation as an apoenzyme referred to as nucleotide-free RAS (nf-RAS).

### ADT-007 inhibits GTP activation of recombinant nf-RAS

We next determined if ADT-007 disrupts RAS-GTP effector binding using recombinant full-length KRAS^WT^. KRAS was treated with EDTA to chelate Mg^+2^, simulating GEF removal of GDP, to generate nf-RAS. ADT-007 treatment of nf-RAS before GTP addition inhibited RAS-RAF binding as determined by RAS-RBD pulldown assays at concentrations comparable to growth IC_50_ values and to an extent approaching the RAS-GDP control (**Figure 3C**). There was a negligible effect of ADT-007 on binding if RAS was preloaded with GTP (**Figure 3D**). A fluorescent GTP analog (Mant-GTP) nucleotide exchange assay was used to determine if ADT-007 disrupts GTP binding to KRAS^WT^. ADT-007 reduced the rate of GTP binding to RAS if pretreated with EDTA to chelate Mg^2+^ (**Figure 3E**), but not after adding non-hydrolysable GTPγS (**Figure 3F**). The pan GTPase inhibitor, CID1067700, inhibited Mant-GTP binding to RAS to the same extent as ADT-007 (not shown) [34].

### ADT-007 binds recombinant nf-RAS

Nuclear magnetic resonance (NMR) spectroscopy was used to confirm binding of ADT-007 to recombinant full-length KRAS^WT^. These experiments were conducted by chelating Mg^2+^ with EDTA to disassociate GDP and generate nf-RAS, producing a sample with multiple NMR spectral differences from GDP-KRAS (**Figure 4A**). Removal of GDP included attenuated signals for G13, a hotspot mutation site in the P-loop [35], and K16 (adjacent to S17) critical for Mg^2+^ binding [36], which could be readily assigned due to their unique resonance frequencies. Treatment of nf-RAS with ADT-007 resulted in several discrete changes in HN signals, including attenuated signals from G13 and K16 residues (**Figure 4B**). Consistent with biochemical experiments, the NMR spectra of GDP-KRAS^WT^ in the presence of Mg^2+^ showed no significant differences by ADT-007 treatment, indicating a low likelihood of ADT-007 binding to nucleotide-loaded RAS (not shown).

**Figure 4.**
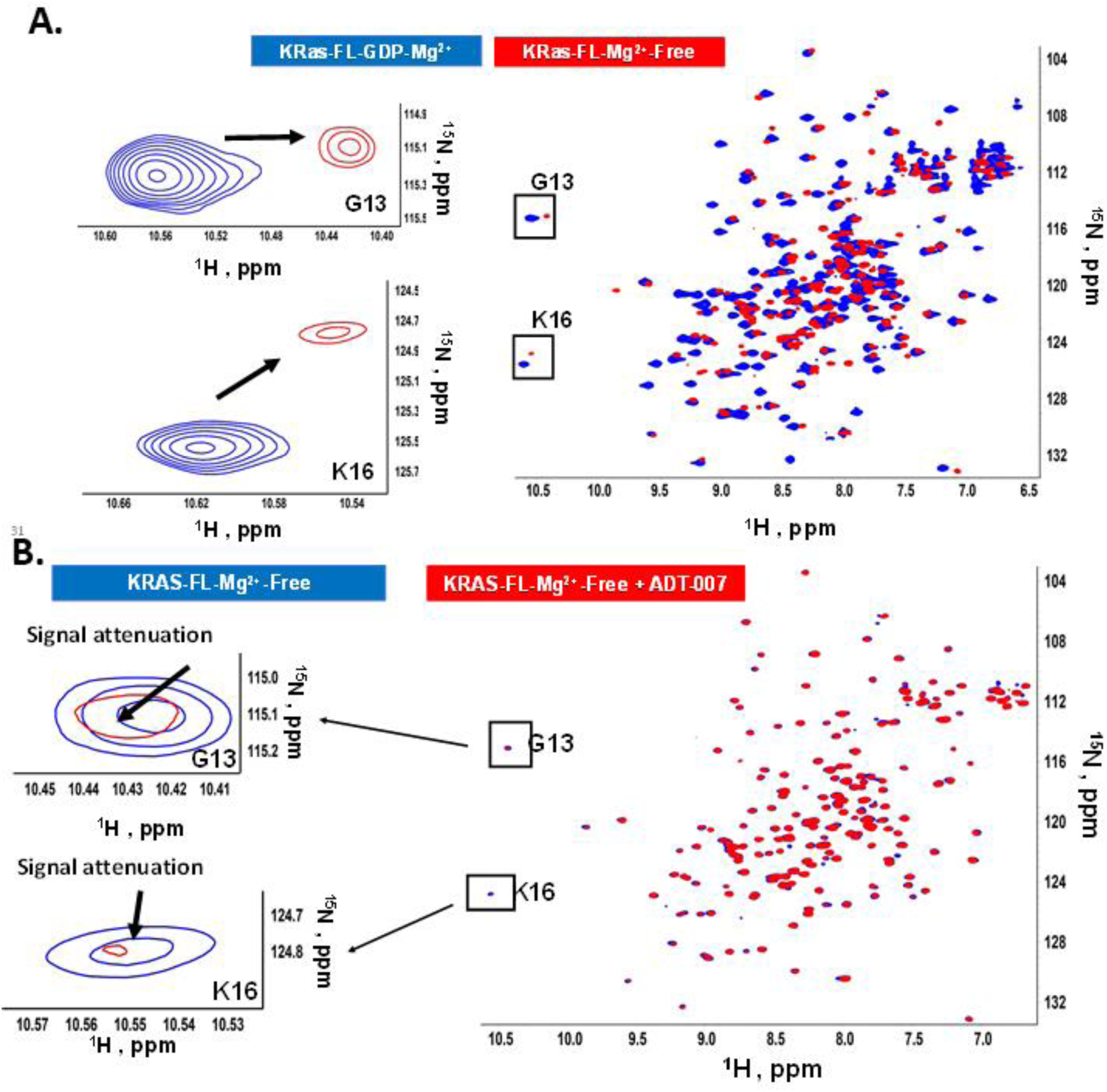
NMR analysis of ADT-007 binding to KRAS. (**A**) NMR spectrum of full length KRAS in the presence (blue) or absence of Mg^2+^ (red) representing nucleotide-loaded or nf-RAS, respectively. (**B**) The addition of ADT-007 (red) to nf-KRAS (blue) (5:1 ratio) resulted in statistically significant NMR signal attenuation at G13 and K16 and other residues (described in methods). Assignments for G13 and K16, due to their unique resonance frequencies, are shown to the left.

### ADT-007 inhibits tumor growth and RAS signaling *in vivo*

As described above, *ex vivo* treatment of mouse liver microsome with ADT-007 identified a metabolic vulnerability of ADT-007 to be rapidly metabolized by glucuronidation (**Figure 2C-D**) prompting experiments to evaluate the antitumor activity of ADT-007 by localized delivery to minimize liver glucuronidation. Localized delivery (peri-tumorally) of ADT-007 completely arrested tumor growth and induced partial regression in the KRAS^G12V^ gall bladder adenocarcinoma PDX model (**Figure 5A**) and reduced tumor growth in a KRAS^G12C^ PDA PDX model (not shown). Intraperitoneal injection of ADT-007 significantly reduced growth in three peritoneal metastasis PDX models bearing mutant KRAS^G12A^ (PMCA-1) (**Figure 5B**), KRAS^G12D^ (PMP1) (**Figure 5C**), or KRAS^WT^ and PDGFR^T922I^ (PC244) (**Figure 5D**). PMCA-1 was the most sensitive to ADT-007 with the growth inhibition persisting long after treatment discontinuation, and two mice tumor-free on day 100. Consistent with *in vitro* experiments described above showing that RAS^WT^ cancer cell lines with upstream mutations are sensitive to ADT-007 (**Table 1**), the PC244 PDX model expressing RAS^WT^ isozymes, but a suspected activating mutation in PDGFR, was sensitive to ADT-007 *in vivo*. Reductions in mucin production were notable by MRI (**Supplemental Figure 6A**) and tumor volume measurements in each model (**Supplemental Figure 6B-D**).

**Figure 5.**
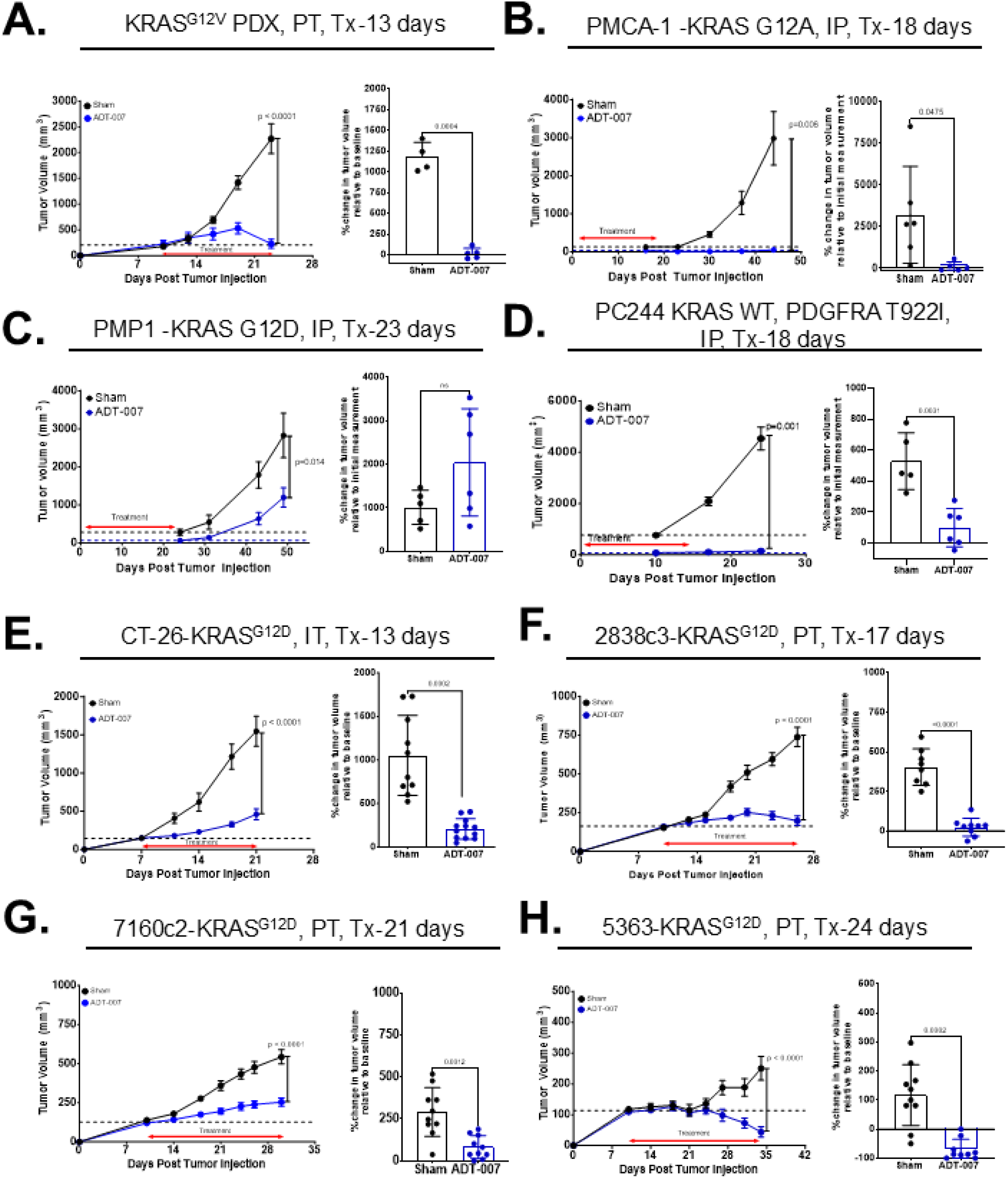
ADT-007 induces growth arrest and regression in mouse tumor models. (**A**) Left panel: Growth inhibition of SQ KRAS^G12V^ PDX gall bladder adenocarcinoma (PT 5 mg/kg, BID, 13 days, n = 5 mice/group, mean ± SEM). Right panel: % change in tumor volume. (**B**-**D**) Left panels: Growth inhibition of mucinous PDX colorectal adenocarcinomas (**B**) PMCA-1, (**C**) PMP1, and (**D**) PC244 in nude mice (IP, 2.5 mg/kg, BID, treatment periods as indicated, n = 5-6 mice/group, mean ± SEM). Right panels: % change in tumor volume. (**E**) Left panel: Growth inhibition of SQ CT26 CRC tumors (IT, 10 mg/kg, QD, 13 days, mean ± SEM, n=12). Right panel: % change in tumor volume. (**F-H**) Left panels: Growth inhibition of SQ (**F**) 2838c3, (**G**) 7160c2, (**H**) 5363 PDA tumors in C57BL/6 mice (PT, 5 mg/kg, BID, treatment periods as indicated) (n = 7 - 10 mice/group, mean ± SEM). Right Panels: % change in tumor volume. All animal studies are representative of 2-3 independent experiments. Left panels: Statistical significance was assessed using 2-way ANOVA with Sadik’s multiple comparisons test. Right panels: Statistical significance was assessed with Welch’s t-test.

The antitumor activity of ADT-007 was also determined against an array of syngeneic CRC and PDA tumor models bearing KRAS^G12D^ mutations grown SQ in immune-competent mice. When administered locally by intra-tumor (IT) injection, ADT-007 (10 mg/kg, QD) strongly inhibited SQ tumor growth (T/C_max_ of 27%) in the KRAS mutant CT26 colon tumor model (**Figure 5E, left panel**) with an 80% reduction in mean tumor volume at the end of the treatment compared to sham-treated mice (**Figure 5E, right panel**). The extent of tumor growth inhibition by ADT-007 in each mouse was closely associated with the extent of reduction of phosphorylated RAF, MEK, and ERK (**Supplemental Figure 7**). Peritumoral administration of ADT-007 (5 mg/kg, BID) induced growth arrest in previously characterized syngeneic mouse PDA cell lines 2838c3 (**Figure 5F**) and 7160c2 (**Figure 5G**) [25], and tumor regressions in a KPC-derived PDA cell line 5363 (**Figure 5H**).

Although ADT-007 was rapidly metabolized by the liver, the compound was stable in mouse plasma (**Supplemental Figure 8A**). We therefore synthesized a carbamate prodrug of ADT-007, ADT-316, to protect the vulnerable hydroxyl group from glucuronidation (**Supplemental Figure 8B**). Oral administration of ADT-316 generated plasma levels of ADT-007 appreciably higher than growth inhibitory IC_50_ values for at least 8 h following a single dose (**Supplemental Figure 8C**). While PK analysis of plasma levels of ADT-316 and ADT-007 revealed modest conversion of the prodrug to ADT-007, ADT-316 significantly inhibited tumor growth in the CT26 murine CRC model at a well-tolerated dosage (100 mg/kg, bid) (**Supplemental Figure 8D**). Given that the incomplete conversion of ADT-316 to ADT-007 likely accounts for its modest antitumor activity, these results support further development of prodrugs to generate higher or more sustained systemic ADT-007 levels.

### ADT-007-mediated inhibition of KRAS^G12D^ alters the tumor immune microenvironment (TiME)

To determine the impact of ADT-007 on tumor immunity in mutant KRAS^G12D^ tumors, we assessed the activity of ADT-007 in C57BL/6 recipient mice using 2 PDA tumor lines bearing KRAS^G12D^. Bot cell lines (2838c3 and 7160c2) has been previously characterized as tumor models with increased T cell infiltrate, yet exhibiting different abundances of myeloid subsets (**Supplemental Figure 9A-B**), basal levels of T cell activation, and function [25]. At treatment termination, ADT-007 decreased the proportion of YFP^+^ tumor cells in both models (**Figures 6A and 6I**). ADT-007 treatment increased proportions of gMDSC and mMDSC subsets in 7160c2 (**Figure 6B**), but not in 2838c3 tumor-bearing mice, which exhibited a significant decrease in macrophages (**Figure 6J**) in response to treatment. No changes in dendritic cell subset numbers were noted in either model (**Figures 6C***,7160c2* and **6K***,2838c3*). Analysis of macrophage phenotypes in both models revealed trends for decreasing MHCII and CD206 expression (**Supplemental Figures 9C***,2838c3* and **9D**,7160c2).

**Figure 6.**
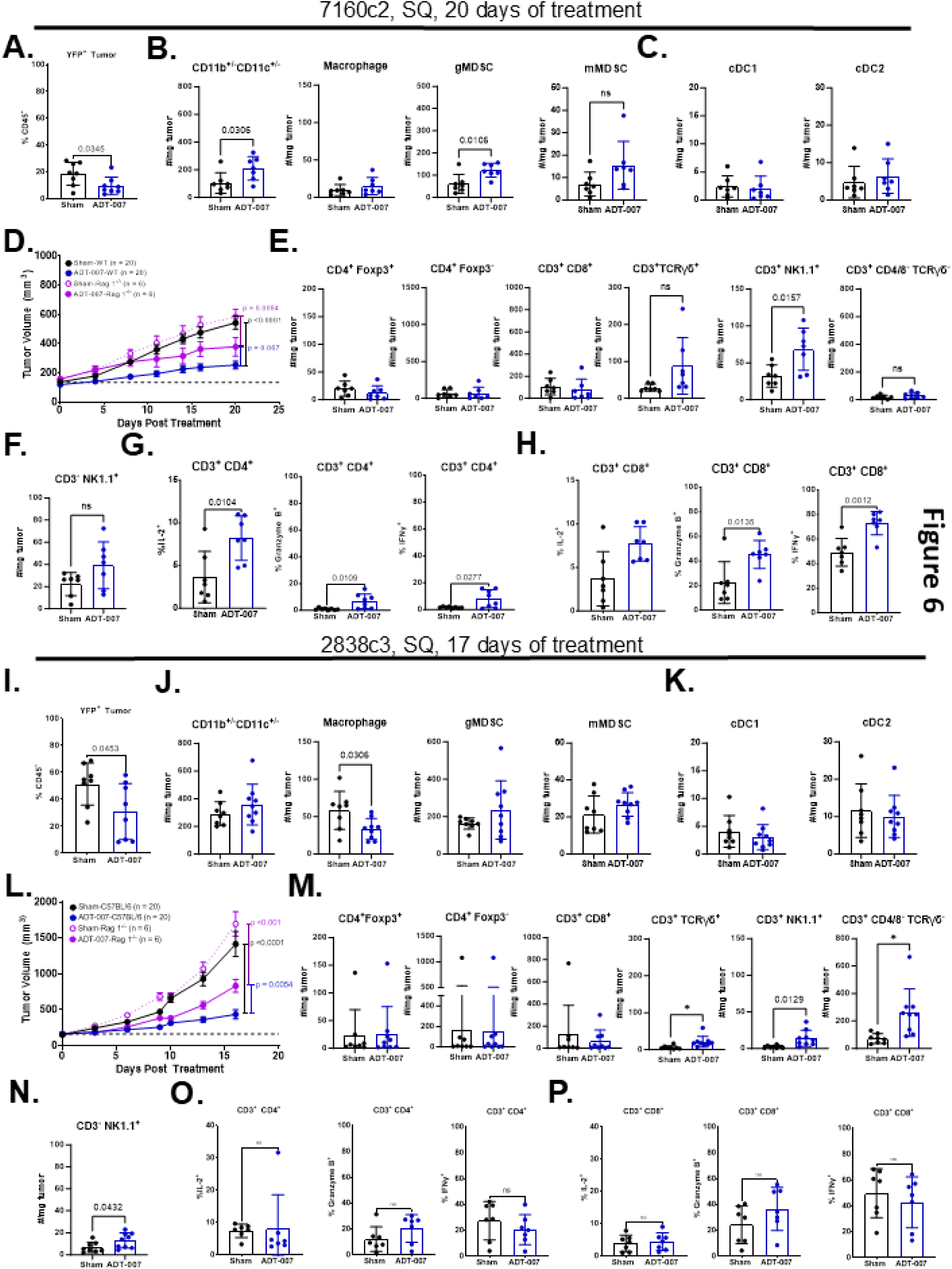
ADT-007 modulates tumor immunity in the PDA TME. **(A-H)** <u>7160c2 PDA model (20 day of treatment) and</u> (**I-P**) <u>2838c3 PDA model (17 day of treatment).</u> (**A and I**) % CD45^-^ YFP^+^ tumor cells in ADT-007 vs. sham (n = 8). (**B and J**) Numbers/mg tumor of total myeloid (CD11b^+/-^CD11c^+/-^, macrophage, gMDSC, and mMDSC (**Supplemental Figure 8A**-*gating scheme*) in ADT-007 treated vs. sham (n = 8). (**C and** Numbers/mg tumor of cDC1 and cDC2 subsets in ADT-007 vs. sham (n = 8). (**D and** Tumor growth in 7160c2 PDA tumor-bearing C57BL/6 WT vs. C57BL/6 Rag 1^-/-^ mice (SQ, 5 mg/kg, BID, days 7-28, n = 6-22 mice/group, mean ± SEM). (**E and M**) Numbers/mg tumor of CD4^+^ Foxp3^+^, CD4^+^ Foxp3^-^, CD8^+^ T, γδ^+^ (TCRγδ^+^), NK T (CD3^+^ NK1.1^+^), and CD4/CD8^-^ / γδ^-^ T cells and (**F and N**) NK (CD3^-^ NK1.1^+^) cells (**Supplemental Figure 9A**-*gating scheme*) in ADT-007 vs. sham (n = 8). (**G-H and O-P**) % intracellular cytokine (IL-2, granzyme B, and IFNγ) production (ICS) from (**G and O**) CD4^+^ and (**H and P**) CD8^+^ T cells were evaluated after 5-h stimulation with PMA/ionomycin + brefeldin A in ADT-007 vs. sham (n = 8). Column graphs - 2 independent experiments, statistical significance: Welch’s t-test, error bars - SD. Tumor growth curves - aggregate of 2 independent experiments, statistical significance: 2-way ANOVA with Tukey’s multiple comparison, error bars - SEM.

Efficacy ADT-007 was partially dependent upon T and B cells, as its efficacy was reduced in immune-deficient C57BL/6 Rag 1^-/-^ *vs*. C57BL/6 mice (**Figures 6D***,7160c2* and **6L***,2838c3*). Both models contained similar numbers of T cells prior to treatment (**Supplemental Figure 10A-B**). ADT-007 treatment increased numbers of γδ, NKT, and NK cells, whereas CD4^+^ and CD8^+^ T cell numbers remained unchanged within the TiME in both models (**Figures 6E-F***,7160c2* **and 6M-N***,2838c3*). The activation status of T cell subsets (**Supplemental Figure 10A**) in the 7160c2, but not 2838c3 TiME, shifted to a CD44^hi^ CD62L^-^effector phenotype with ADT-007 treatment (**Supplemental Figure 10C-D)** and PD-1 increased on CD4^+^ and CD8^+^ T cells in 7610c2, but not 2838c3 TiME (**Supplemental Figure 11A**,7160c2 and **11B***,2838c3*), despite higher baseline levels of PD-1 expression (**Supplemental Figure 11C**). Co-expression of PD-1 and TIM-3 on tumor infiltrating T cells was modestly increased only in 7160c2 model after ADT-007 treatment (**Supplemental Figure 11D-E**).

In the 7160c2 TiME both CD4^+^ and CD8^+^ T cells exhibited increased expression of IL-2, granzyme B, and IFNγ (**Figure 6G-H**) after stimulation with PMA/ionomycin, whereas minimal changes in cytokine expression were noted in T cells within the 2838c3 TiME (**Figure 6O-P**). Using an IFNγ-Thy1.1 reporter model system [24], we assessed IFNγ production marked by surface Thy1.1 on innate and adaptive T and NK subsets. Consistent with intracellular cytokine staining (ICS) results, ADT-007 increased IFNγ production in CD4^+^ and CD8^+^ T cell subsets in the 7160c2, but not in 2838c3 TiME. In NKT and γδ T cells, IFNγ was unchanged (**Supplemental Figure 12A***,7160c2* **and 12B***,2838c3*), whereas a decrease was noted in NK cells from ADT-007 treated tumors (**Supplemental Figure 12A***,7160c2* **and 12B***,2838c3*).

## DISCUSSION

ADT-007 potently inhibited the growth of cancer cells with mutant RAS regardless of the RAS mutation or isozyme. Cancer cells with RAS^WT^ but mutations resulting in constitutive activation of RTK, loss-of-function in GAPs were comparably sensitive to ADT-007 as was a gastric cancer cell line harboring gene amplification of the KRAS^WT^ gene. Despite broad activity, ADT-007 displayed exquisite selectivity as cancer cells with BRAF mutations, immediately downstream from RAS, and cells from normal tissues were essentially insensitive to ADT-007. ADT-007 sensitivity was attributed to high RAS-GTP level and dependence on RAS signaling for proliferation (e.g., RAS addiction), while normal cells or cancer cell lines with RAS^WT^ have a metabolic mechanism protecting them from ADT-007 cytotoxicity involving glucuronidation.

Concentrations of ADT-007 effective for binding RAS in cells or recombinant RAS were within the same range as concentrations required to inhibit growth, colony formation, and induce mitotic arrest and apoptosis, as well as concentrations that block RAS activation and MAPK/AKT signaling. The inhibitory effect of ADT-007 on RAS-GTP occurred by treating KRAS mutant cancer cells under standard growth conditions, serum-free conditions, or following EGF stimulation. ADT-007 reduced RAS-GTP from mutant KRAS and co-expressed HRAS^WT^ and NRAS^WT^ isozymes that were activated by EGF. This activity may be a key advantage over mutant-specific KRAS or pan-KRAS inhibitors to circumvent resistance from compensatory activity of unchecked RAS^WT^ NRAS or HRAS isozymes, although as discussed below ADT-007 may have additional advantages such as its unique ability to induce mitotic arrest and apoptosis that may also avert resistance.

Localized delivery of ADT-007 strongly inhibited tumor growth in immunodeficient and syngeneic murine models of CRC and PDA with partial dependence on the adaptive immune system. ADT-007 activated T cells and enhanced their cytotoxic functions in KRAS^G12D^ mutated CRC and PDA, immune-competent mouse models. Oral delivery of a prototypic prodrug, ADT-316, significantly inhibited tumor growth but with modest activity likely because of incomplete conversion to ADT-007. Recognizing the limitations of ADT-316, we designed an improved series of prodrugs that are currently under study, one of which, ADT-1004, shows robust antitumor activity and exquisite RAS selectivity in mouse models involving mouse or human PDA cell lines or PDX models of PDA. ADT-1004 also shows superior antitumor activity over sotorasib and adagrasib in experiments comparing equal dosages of each compound administer orally in mice implanted with resistant MIA PaCa-2 PDA cells (bioRxiv 2024. 10.1101/2024.10.04.616725).

ADT-007 appears to bind RAS within the nucleotide binding domain to block GTP-activation of effector interactions and downstream MAPK/AKT signaling. Molecular modeling studies provided initial evidence that ADT-007 binds RAS within the nucleotide-binding domain, which were supported by biochemical experiments showing that ADT-007 inhibits GTP binding to recombinant RAS and GTP-dependent RAS-RAF interactions. These effects were only observed under nucleotide-free conditions, but not under nucleotide loading conditions. The selectivity of ADT-007 to bind nf-RAS was confirmed by NMR spectroscopy experiments, which provided additional evidence for binding within the nucleotide-binding domain, although further studies are needed to define the mechanism of binding more precisely.

ADT-007 binding to nf-RAS could negatively impact its antitumor activity, given this is a transitional conformation and the presumed temporal nature of target exposure. While *in vitro* experiments show near complete killing of RAS mutant cancer cells by ADT-007 given adequate time, an important consideration for *in vivo* antitumor activity is if ADT-007 has a sufficiently long plasma half-life for target occupancy. Indeed, oral administration of ADT-316 and ADT-1004 generated sustained plasma levels of ADT-007 in mice that were appreciably greater than *in vitro* growth IC_50_ values and significantly inhibited tumor growth *in vivo*. In addition, previous reports that a significant percentage of RAS exists in cells as an apoenzyme and that a nf-RAS monobody inhibits tumor growth [37], suggest that ADT-007 binding to nf-RAS can overcome this perceived liability.

Metabolic deactivation of ADT-007 by UGT-mediated glucuronidation presents implications for the interpretation of both *in vitro* and *in vivo* experiments. Liver microsome experiments showed that ADT-007 is rapidly glucuronidated by UGT enzymes ubiquitously expressed in normal tissues and well known to metabolize compounds with phenolic hydroxyl moieties. Unexpectedly, RAS^WT^ cancer cells with mutant BRAF had high levels of major UGT isozymes commonly expressed in normal cells. In contrast, cancer cells with mutant KRAS had low levels of UGT isozymes. Consistent with these differences, low ADT-007 levels and glucuronide-conjugated ADT-007 were measured in RAS^WT^ cells, while high ADT-007 levels but no glucuronide-conjugated ADT-007 were measured in KRAS mutant HCT-116 cells by LC-MS/MS. Moreover, conditioned medium obtained from RAS^WT^ cells exposed to ADT-007, but not KRAS mutant cells, completely lost growth inhibitory activity, reflecting metabolic deactivation of ADT-007. Pretreatment with a known competitive UGT inhibitor, propofol, also increased sensitivity of RAS^WT^ cells to ADT-007 but did not significantly impact sensitivity of mutant KRAS cells to ADT-007. These results suggest that UGT-mediated glucuronidation protects bystander cells from normal tissues from pan-RAS inhibition by ADT-007. Despite acting as a pan-RAS inhibitor, which otherwise would be considered lethal based on genomic knock-out experiments in mice [38, 39], ADT-007 has a unique metabolic mechanism of RAS selectivity. While UGT isozymes are well known to be expressed in normal tissues, including blood cells, and regulated by multiple signaling pathways and transcription factors [40], further study is needed to understand how UGT expression is suppressed by RAS mutations.

Other sulindac derivatives were previously reported to inhibit cancer cell growth by a dual mechanism involving inhibition of cyclic guanine monophosphate phosphodiesterase (cGMP PDE) activity and tubulin polymerization [31]. The former was based on reports that derivatives with a trimethoxy substitution inhibited cGMP PDE activity at concentrations that inhibit cancer cell growth [21, 41], while the latter was based on their chemical resemblance with colchicine, which is known to bind tubulin and disrupt microtubules [42]. We, therefore, determined if ADT-007 inhibits cGMP PDE activity by measuring effects on cGMP hydrolysis using recombinant PDE10, a cGMP-degrading PDE isozyme (not shown), or capacity to bind PDE10 in cells using cellular thermal stability assays **(Figure 3A)**, but found no evidence of PDE10 inhibition or binding, respectively. In addition, ADT-007 only inhibited tubulin polymerization at concentrations ∼100-fold higher than growth IC_50_ values (**Supplemental Figure 3H**). These observations lead us to conclude that neither cGMP PDE nor tubulin polymerization inhibition account for the growth inhibitory activity of ADT-007. Reports that tubulin inhibitors induce MAPK pathway activity [43], rather than inhibit MAPK signaling, provide additional evidence that ADT-007 acts by targeting RAS. Moreover, colchicine lacked selectivity in growth assays involving RAS mutant vs. RAS^WT^ cancer cells (not shown). We also tested the topoisomerase inhibitor, etoposide, because it has the same phenolic hydroxyl moiety as ADT-007, which we show is essential for RAS binding but found no evidence of RAS selectivity in growth assays.

Unlike KRAS^G12C^ inhibitors that have limited use for patients with this relatively rare mutation [30, 44], ADT-007 has broad anticancer activity with potential to be developed as a generalizable monotherapy for the majority of patients with RAS mutant cancers. In addition, a significant proportion of patients with RTK fusions that hyper-activate RAS^WT^ isozymes [45] would be expected to be sensitive to ADT-007. Evidence indicates that ADT-007 exhibits high potency in tumor cells with activated RAS signaling, which may provide greater therapeutic utility in patients with metastatic disease in which the proliferation of cancer cells depends upon activated KRAS signaling for survival in the metastatic niche [46].

Equally significant to the broad antitumor activity of ADT-007 is its potential to provide a second-line therapy for patients with KRAS mutant cancers that develop resistance to other RAS inhibitors, for example, from co-occurring RAS mutations or compensation from RAS^WT^ isozymes [7, 20]. We show here that ADT-007 maintains growth inhibitory activity in LLC cells with co-occurring KRAS^G12C^ and NRAS^Q61H^ mutations [47], while the KRAS^G12C^ inhibitor, sotorasib, and the pan-KRAS inhibitor, BI-2865 lost growth inhibitory activity. Other experiments showed that ADT-007 induces mitotic arrest and apoptosis resulting in cytotoxicity, while BI-2865 or AMG-510 did not, and another pan-RAS inhibitor, RMC-2865, did not induce apoptosis, appearing to be limited to a cytostatic mechanism. This difference was apparent in clonogenic colony formation assays in which ADT-007 treatment caused near complete inhibition and to a much greater extent than BI-2865 or RMC-6236 when compared side-by-side. These advantages of ADT-007 over other RAS inhibitors as observed *in vitro* awaits further *in vivo* studies but suggests the potential to escape intrinsic and acquired resistance.

ADT-007-mediated *in vivo* tumor growth inhibition also appears to be partially dependent upon the adaptive immune response as evidenced by reduced antitumor activity of ADT-007 in 2838c3 and 7160c2 tumors grown in immune-deficient Rag 1^-/-^ mice lacking B and T cells *vs*. in immune-competent C57BL/6. Similar findings were observed with KRAS^G12C^ [48, 49] and KRAS^G12D^-specific inhibitors [50], demonstrating that T cells contribute to the efficacy of other RAS inhibitors. The presence of T cells predicted synergy with PD-1 immune checkpoint blockade in preclinical models of NSCLC and PDA [49, 51]. However, the importance of B cells to anti-tumor immunity of ADT-007 requires further study.

In T cell inflamed PDA tumor models (2838c3 and 7160c2), ADT-007 treatment increased numbers of innate-like T lymphocytes (NK T (CD3^+^ NK1.1^+^), γδT, and CD3^+^ CD4^-^/CD8^-^ T, and NK (CD3^-^ NK1.1^+^) cells), whereas conventional CD4^+^ and CD8^+^ T cell numbers did not change. Analysis of IFNγ levels using Thy1.1-IFNγ reporter mice revealed that most adaptive and innate-like T cells were Thy1.1^+^, consistent with high levels of IFNγ production *in vivo*, whereas, differences in ADT-007 on T cell activation and cytokine secretion in both models, may be explained by the higher baseline activation and cytokine secretion of T cells isolated from the 2838c3 model system [25], suggesting that differences in baseline T cell activation and function may contribute to differences in outcomes with ADT-007 treatment in different tumor models. Overall, these findings suggest role(s) for innate and adaptive immune response in the antitumor activity of ADT-007 that can potentially be exploited with relevant immune therapy combinations.

In conclusion, ADT-007 represents a 1^st^-in-class pan-RAS inhibitor with potential to address the complex RAS mutational landscape of many human cancers and properties superior to other RAS inhibitors to circumvent resistance. These results support IND-enabling studies of an ADT-007 analog or prodrug as a generalizable monotherapy or combined with other RAS pathway-directed inhibitors or immunotherapy for treating RAS-driven cancers.

## References

1. Siegel RL, Giaquinto AN, Jemal A: Cancer statistics, 2024. CA Cancer J Clin 2024, 74(1):12-49.

2. Yang Y, Zhang H, Huang S, Chu Q: KRAS Mutations in Solid Tumors: Characteristics, Current Therapeutic Strategy, and Potential Treatment Exploration. J Clin Med 2023, 12(2).

3. Hingorani SR, Petricoin EF, Maitra A, Rajapakse V, King C, Jacobetz MA, Ross S, Conrads TP, Veenstra TD, Hitt BA et al: Preinvasive and invasive ductal pancreatic cancer and its early detection in the mouse. Cancer Cell 2003, 4(6):437–450.

4. Kojima K, Vickers SM, Adsay NV, Jhala NC, Kim HG, Schoeb TR, Grizzle WE, Klug CA: Inactivation of Smad4 accelerates Kras(G12D)-mediated pancreatic neoplasia. Cancer Res 2007, 67(17):8121–8130.

5. Boutin AT, Liao WT, Wang M, Hwang SS, Karpinets TV, Cheung H, Chu GC, Jiang S, Hu J, Chang K et al: Oncogenic Kras drives invasion and maintains metastases in colorectal cancer. Genes Dev 2017, 31(4):370–382.

6. Bardeesy N, Aguirre AJ, Chu GC, Cheng KH, Lopez LV, Hezel AF, Feng B, Brennan C, Weissleder R, Mahmood U et al: Both p16(Ink4a) and the p19(Arf)-p53 pathway constrain progression of pancreatic adenocarcinoma in the mouse. Proc Natl Acad Sci U S A 2006, 103(15):5947–5952.

7. Sheffels E, Kortum RL: The Role of Wild-Type RAS in Oncogenic RAS Transformation. Genes (Basel) 2021, 12(5).

8. Ying H, Kimmelman AC, Lyssiotis CA, Hua S, Chu GC, Fletcher-Sananikone E, Locasale JW, Son J, Zhang H, Coloff JL et al: Oncogenic Kras maintains pancreatic tumors through regulation of anabolic glucose metabolism. Cell 2012, 149(3):656–670.

9. Mukhopadhyay S, Goswami D, Adiseshaiah PP, Burgan W, Yi M, Guerin TM, Kozlov SV, Nissley DV, McCormick F: Undermining Glutaminolysis Bolsters Chemotherapy While NRF2 Promotes Chemoresistance in KRAS-Driven Pancreatic Cancers. Cancer Res 2020, 80(8):1630–1643.

10. Collins MA, Brisset JC, Zhang Y, Bednar F, Pierre J, Heist KA, Galban CJ, Galban S, di Magliano MP: Metastatic pancreatic cancer is dependent on oncogenic Kras in mice. PLoS One 2012, 7(12):e49707.

11. Zhang Y, Yan W, Collins MA, Bednar F, Rakshit S, Zetter BR, Stanger BZ, Chung I, Rhim AD, di Magliano MP: Interleukin-6 is required for pancreatic cancer progression by promoting MAPK signaling activation and oxidative stress resistance. Cancer Res 2013, 73(20):6359–6374.

12. Zdanov S, Mandapathil M, Abu Eid R, Adamson-Fadeyi S, Wilson W, Qian J, Carnie A, Tarasova N, Mkrtichyan M, Berzofsky JA et al: Mutant KRAS Conversion of Conventional T Cells into Regulatory T Cells. Cancer Immunol Res 2016, 4(4):354–365.

13. Bayne LJ, Beatty GL, Jhala N, Clark CE, Rhim AD, Stanger BZ, Vonderheide RH: Tumor-derived granulocyte-macrophage colony-stimulating factor regulates myeloid inflammation and T cell immunity in pancreatic cancer. Cancer Cell 2012, 21(6):822–835.

14. Ischenko I, D’Amico S, Rao M, Li J, Hayman MJ, Powers S, Petrenko O, Reich NC: KRAS drives immune evasion in a genetic model of pancreatic cancer. Nat Commun 2021, 12(1):1482.

15. Coelho MA, de Carne Trecesson S, Rana S, Zecchin D, Moore C, Molina-Arcas M, East P, Spencer-Dene B, Nye E, Barnouin K et al: Oncogenic RAS Signaling Promotes Tumor Immunoresistance by Stabilizing PD-L1 mRNA. Immunity 2017, 47(6):1083–1099 e1086.

16. Muthalagu N, Monteverde T, Raffo-Iraolagoitia X, Wiesheu R, Whyte D, Hedley A, Laing S, Kruspig B, Upstill-Goddard R, Shaw R et al: Repression of the Type I Interferon Pathway Underlies MYC- and KRAS-Dependent Evasion of NK and B Cells in Pancreatic Ductal Adenocarcinoma. Cancer Discov 2020, 10(6):872–887.

17. Stalnecker CA, Der CJ: RAS, wanted dead or alive: Advances in targeting RAS mutant cancers. Sci Signal 2020, 13(624).

18. Kwan AK, Piazza GA, Keeton AB, Leite CA: The path to the clinic: a comprehensive review on direct KRAS(G12C) inhibitors. J Exp Clin Cancer Res 2022, 41(1):27.

19. Ryan MB, Fece de la Cruz F, Phat S, Myers DT, Wong E, Shahzade HA, Hong CB, Corcoran RB: Vertical Pathway Inhibition Overcomes Adaptive Feedback Resistance to KRAS(G12C) Inhibition. Clin Cancer Res 2020, 26(7):1633–1643.

20. Awad MM, Liu S, Rybkin, II, Arbour KC, Dilly J, Zhu VW, Johnson ML, Heist RS, Patil T, Riely GJ et al: Acquired Resistance to KRAS(G12C) Inhibition in Cancer. N Engl J Med 2021, 384(25):2382–2393.

21. Thompson HJ, Jiang C, Lu J, Mehta RG, Piazza GA, Paranka NS, Pamukcu R, Ahnen DJ: Sulfone metabolite of sulindac inhibits mammary carcinogenesis. Cancer Res 1997, 57(2):267–271.

22. Herrmann C, Block C, Geisen C, Haas K, Weber C, Winde G, Moroy T, Muller O: Sulindac sulfide inhibits Ras signaling. Oncogene 1998, 17(14):1769–1776.

23. Karaguni IM, Glusenkamp KH, Langerak A, Geisen C, Ullrich V, Winde G, Moroy T, Muller O: New indene-derivatives with anti-proliferative properties. Bioorg Med Chem Lett 2002, 12(4):709–713.

24. Harrington LE, Janowski KM, Oliver JR, Zajac AJ, Weaver CT: Memory CD4 T cells emerge from effector T-cell progenitors. Nature 2008, 452(7185):356-360.

25. Li J, Byrne KT, Yan F, Yamazoe T, Chen Z, Baslan T, Richman LP, Lin JH, Sun YH, Rech AJ et al: Tumor Cell-Intrinsic Factors Underlie Heterogeneity of Immune Cell Infiltration and Response to Immunotherapy. Immunity 2018, 49(1):178–193 e177.

26. Flatmark K, Reed W, Halvorsen T, Sorensen O, Wiig JN, Larsen SG, Fodstad O, Giercksky KE: Pseudomyxoma peritonei--two novel orthotopic mouse models portray the PMCA-I histopathologic subtype. BMC Cancer 2007, 7:116.

27. Flatmark K, Davidson B, Kristian A, Stavnes HT, Førsund M, Reed W: Exploring the peritoneal surface malignancy phenotype—a pilot immunohistochemical study of human pseudomyxoma peritonei and derived animal models. Human Pathology 2010, 41(8):1109–1119.

28. Fleten KG, Lund-Andersen C, Waagene S, Abrahamsen TW, Mørch Y, Boye K, Torgunrud A, Flatmark K: Experimental Treatment of Mucinous Peritoneal Metastases Using Patient-Derived Xenograft Models. Transl Oncol 2020, 13(8):100793.

29. Barretina J, Caponigro G, Stransky N, Venkatesan K, Margolin AA, Kim S, Wilson CJ, Lehar J, Kryukov GV, Sonkin D et al: The Cancer Cell Line Encyclopedia enables predictive modelling of anticancer drug sensitivity. Nature 2012, 483(7391):603-607.

30. Waters AM, Der CJ: KRAS: The Critical Driver and Therapeutic Target for Pancreatic Cancer. Cold Spring Harb Perspect Med 2018, 8(9).

31. Xiao D, Deguchi A, Gundersen GG, Oehlen B, Arnold L, Weinstein IB: The sulindac derivatives OSI-461, OSIP486823, and OSIP487703 arrest colon cancer cells in mitosis by causing microtubule depolymerization. Mol Cancer Ther 2006, 5(1):60-67.

32. Lee KJ, Chang WL, Chen X, Valiyaveettil J, Ramirez-Alcantara V, Gavin E, Musiyenko A, Madeira da Silva L, Annamdevula NS, Leavesley SJ et al: Suppression of Colon Tumorigenesis in Mutant Apc Mice by a Novel PDE10 Inhibitor that Reduces Oncogenic β-Catenin. Cancer Prev Res (Phila) 2021, 14(11):995–1008.

33. Hunter JC, Gurbani D, Ficarro SB, Carrasco MA, Lim SM, Choi HG, Xie T, Marto JA, Chen Z, Gray NS et al: In situ selectivity profiling and crystal structure of SML-8-73-1, an active site inhibitor of oncogenic K-Ras G12C. Proc Natl Acad Sci U S A 2014, 111(24):8895–8900.

34. Agola JO, Hong L, Surviladze Z, Ursu O, Waller A, Strouse JJ, Simpson DS, Schroeder CE, Oprea TI, Golden JE et al: A competitive nucleotide binding inhibitor: in vitro characterization of Rab7 GTPase inhibition. ACS Chem Biol 2012, 7(6):1095–1108.

35. Hunter JC, Manandhar A, Carrasco MA, Gurbani D, Gondi S, Westover KD: Biochemical and Structural Analysis of Common Cancer-Associated KRAS Mutations. Mol Cancer Res 2015, 13(9):1325–1335.

36. John J, Rensland H, Schlichting I, Vetter I, Borasio GD, Goody RS, Wittinghofer A: Kinetic and structural analysis of the Mg(2+)-binding site of the guanine nucleotide-binding protein p21H-ras. J Biol Chem 1993, 268(2):923–929.

37. Khan I, Koide A, Zuberi M, Ketavarapu G, Denbaum E, Teng KW, Rhett JM, Spencer-Smith R, Hobbs GA, Camp ER et al: Identification of the nucleotide-free state as a therapeutic vulnerability for inhibition of selected oncogenic RAS mutants. Cell Rep 2022, 38(6):110322.

38. Esteban LM, Vicario-Abejon C, Fernandez-Salguero P, Fernandez-Medarde A, Swaminathan N, Yienger K, Lopez E, Malumbres M, McKay R, Ward JM et al: Targeted genomic disruption of H-ras and N-ras, individually or in combination, reveals the dispensability of both loci for mouse growth and development. Mol Cell Biol 2001, 21(5):1444–1452.

39. Johnson L, Greenbaum D, Cichowski K, Mercer K, Murphy E, Schmitt E, Bronson RT, Umanoff H, Edelmann W, Kucherlapati R et al: K-ras is an essential gene in the mouse with partial functional overlap with N-ras. Genes Dev 1997, 11(19):2468–2481.

40. Allain EP, Rouleau M, Lévesque E, Guillemette C: Emerging roles for UDP-glucuronosyltransferases in drug resistance and cancer progression. Br J Cancer 2020, 122(9):1277–1287.

41. Tinsley HN, Gary BD, Thaiparambil J, Li N, Lu W, Li Y, Maxuitenko YY, Keeton AB, Piazza GA: Colon tumor cell growth-inhibitory activity of sulindac sulfide and other nonsteroidal anti-inflammatory drugs is associated with phosphodiesterase 5 inhibition. Cancer Prev Res (Phila) 2010, 3(10):1303–1313.

42. Lu Y, Chen J, Xiao M, Li W, Miller DD: An overview of tubulin inhibitors that interact with the colchicine binding site. Pharm Res 2012, 29(11):2943–2971.

43. Waters AM, Khatib TO, Papke B, Goodwin CM, Hobbs GA, Diehl JN, Yang R, Edwards AC, Walsh KH, Sulahian R et al: Targeting p130Cas- and microtubule-dependent MYC regulation sensitizes pancreatic cancer to ERK MAPK inhibition. Cell Rep 2021, 35(13):109291.

44. Coley AB, Ward A, Keeton AB, Chen X, Maxuitenko Y, Prakash A, Li F, Foote JB, Buchsbaum DJ, Piazza GA: Pan-RAS inhibitors: Hitting multiple RAS isozymes with one stone. Adv Cancer Res 2022, 153:131–168.

45. Luchini C, Paolino G, Mattiolo P, Piredda ML, Cavaliere A, Gaule M, Melisi D, Salvia R, Malleo G, Shin JI et al: KRAS wild-type pancreatic ductal adenocarcinoma: molecular pathology and therapeutic opportunities. J Exp Clin Cancer Res 2020, 39(1):227.

46. di Magliano MP, Logsdon CD: Roles for KRAS in pancreatic tumor development and progression. Gastroenterology 2013, 144(6):1220–1229.

47. Molina-Arcas M, Moore C, Rana S, van Maldegem F, Mugarza E, Romero-Clavijo P, Herbert E, Horswell S, Li LS, Janes MR et al: Development of combination therapies to maximize the impact of KRAS-G12C inhibitors in lung cancer. Sci Transl Med 2019, 11(510).

48. Canon J, Rex K, Saiki AY, Mohr C, Cooke K, Bagal D, Gaida K, Holt T, Knutson CG, Koppada N et al: The clinical KRAS(G12C) inhibitor AMG 510 drives anti-tumour immunity. Nature 2019, 575(7781):217-223.

49. Mugarza E, van Maldegem F, Boumelha J, Moore C, Rana S, Llorian Sopena M, East P, Ambler R, Anastasiou P, Romero-Clavijo P et al: Therapeutic KRAS(G12C) inhibition drives effective interferon-mediated antitumor immunity in immunogenic lung cancers. Sci Adv 2022, 8(29):eabm8780.

50. Kemp SB, Cheng N, Markosyan N, Sor R, Kim IK, Hallin J, Shoush J, Quinones L, Brown NV, Bassett JB et al: Efficacy of a Small-Molecule Inhibitor of KrasG12D in Immunocompetent Models of Pancreatic Cancer. Cancer Discov 2023, 13(2):298–311.

51. Mahadevan KK, McAndrews KM, LeBleu VS, Yang S, Lyu H, Li B, Sockwell AM, Kirtley ML, Morse SJ, Moreno Diaz BA et al: KRAS(G12D) inhibition reprograms the microenvironment of early and advanced pancreatic cancer to promote FAS-mediated killing by CD8(+) T cells. Cancer Cell 2023, 41(9):1606–1620 e1608.

