## Supplementary material for "A Novel Pan-RAS Inhibitor with a Unique Mechanism of Action Blocks Tumor Growth in Mouse Models of GI Cancer": Figures, Legends, Tables, Methods

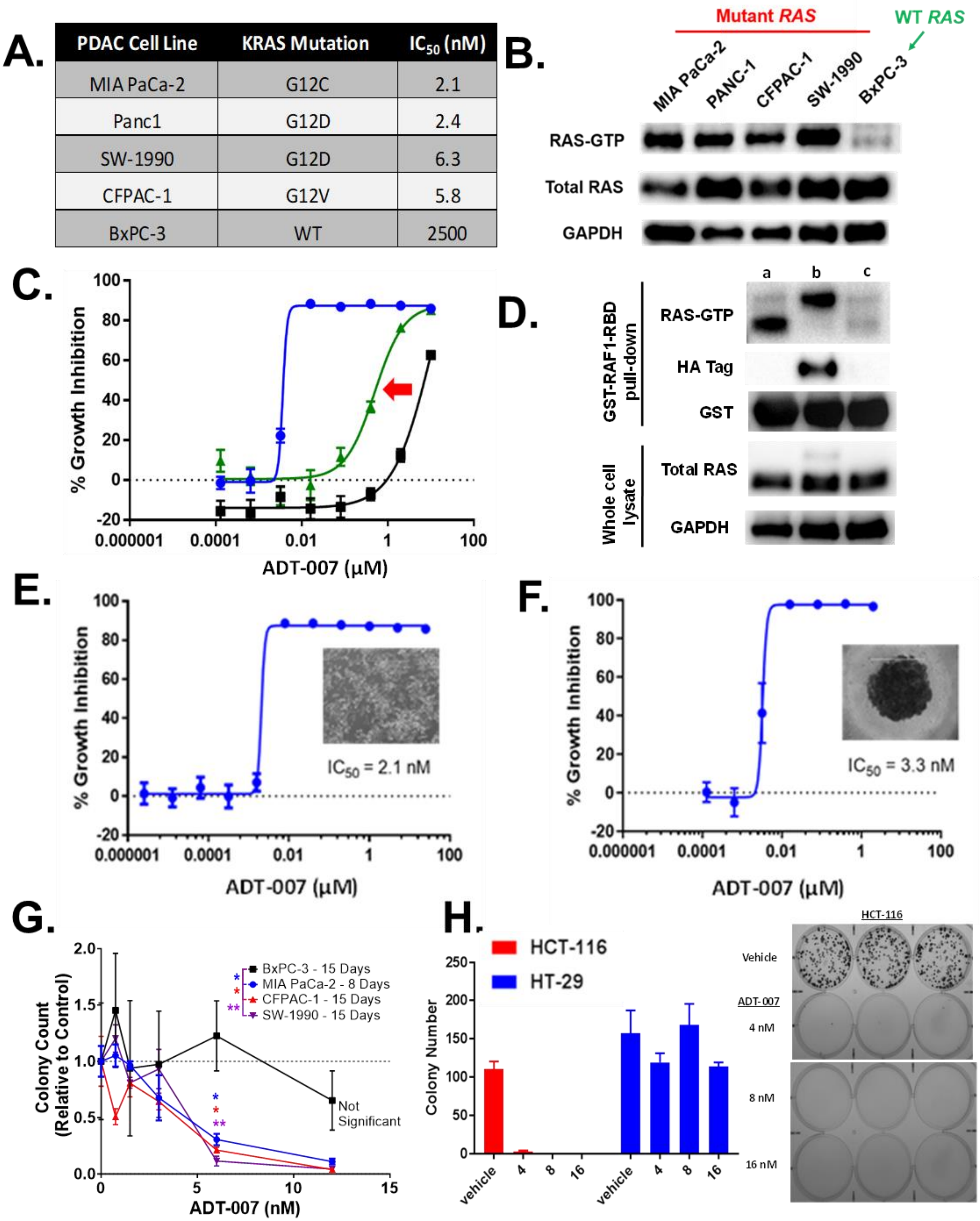


| PDAC Cell Line | KRAS Mutation | $IC_{50}$ (nM) |
|---|---|---|
| MIA PaCa-2 | G12C | 2.1 |
| Panc1 | G12D | 2.4 |
| SW-1990 | G12D | 6.3 |
| CFPAC-1 | G12V | 5.8 |
| BxPC-3 | WT | 2500 |

**Supplemental Figure 1**

**Supplemental Figure 1. ADT-007 potently and selectivity inhibits growth of human RAS mutant pancreatic cancer cells *in vitro*. (A)** Growth inhibition $IC_{50}$ values in MIA PaCa-2, PANC-1, SW-1990, CFPAC-1, and BxPC-3 cells were determined by CTG assay following 96 h of treatment. Data are representative of at least two experiments. KRAS mutations are shown for each cell line. (**B**) Activated RAS (RAS-GTP) levels in PDA cell lines from (**A**) measured by RAS-RBD pulldown assays with total RAS and GAPDH as loading controls. Data are representative of three independent experiments. (**C**) Transfection of $KRAS^{G12C}$ renders $RAS^{WT}$ BxPC-3 cells sensitive to ADT-007 growth inhibition after 72 h of treatment (CTG assay). Parental cells are shown in black, transfected cells shown in green, and MIA PaCa-2 cells as a reference shown in blue. The data are representative of three independent experiments with samples run in triplicate. Error bars represent SEM. (**D**) Activated RAS (RAS-GTP) was measured by RAS-RBD pull-down assays followed by WB using a pan-RAS antibody, HA-Tag and GST antibody with total RAS and GAPDH as loading controls. a, MIA PaCa-2; b, BxPC-3 (HA-$KRAS^{G12C}$); c, BxPC-3 (Parental). (**E-F**) Growth inhibitory activity of ADT-007 in (**E**) a monolayer culture or (**F**) spheroids of MIA PaCa-2 cells as measured by CTG assay. Data are representative of three experiments; error bars represent SEM. (**G**) ADT-007 selective inhibition of colony formation of PDA cell lines with mutant RAS. (Representative of two independent experiments; one-way ANOVA; *$p$<0.05, **p<0.01 *vs*. vehicle (DMSO) control for each indicated cell line). Curves are graphed as the average of 3 replicates ± SEM. (**H**) ADT-007 inhibition of colony formation of $KRAS^{G12D}$ mutant HCT-116 cells without affecting $RAS^{WT}$ HT-29 cells. Bars are graphed as the average of 3 replicates ± SEM. Results shown are representative of at least 2 independent experiments.

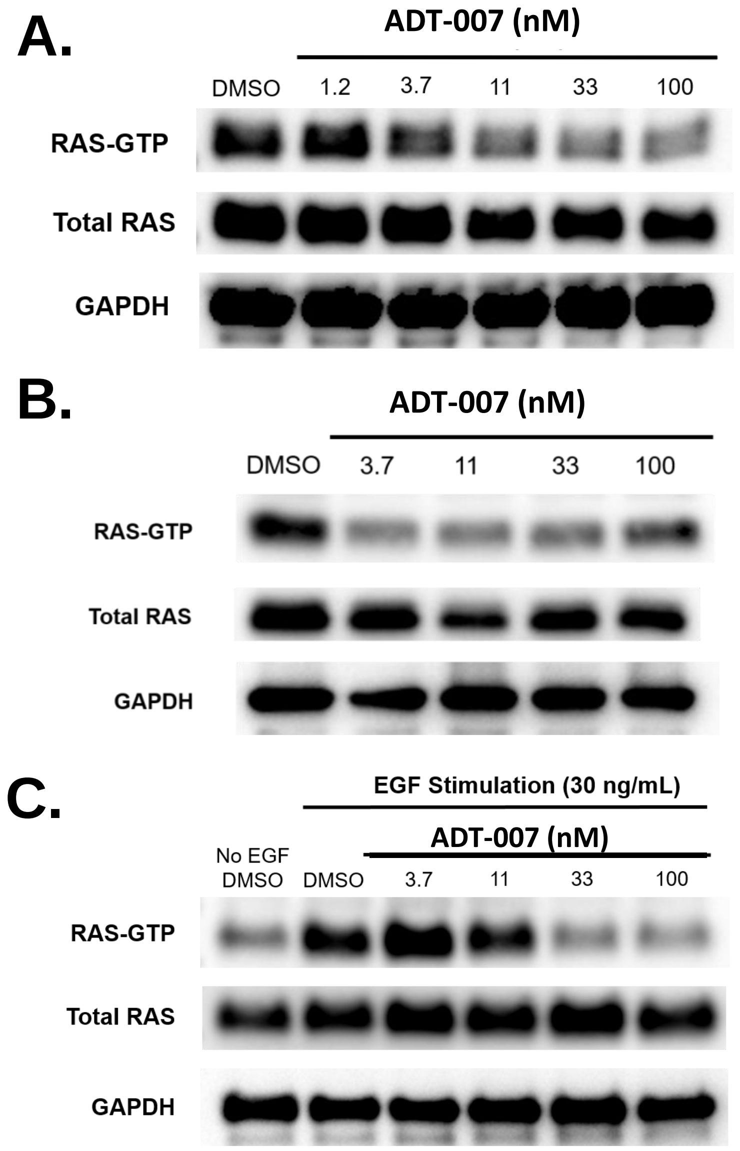


**Supplemental Figure 2**

**Supplemental Figure 2. ADT-007 inhibition of activated RAS.** (**A-C**) ADT-007 inhibits activated RAS (RAS-GTP) in $KRAS^{G12C}$ mutant MIA PaCa-2 PDA cells grown under conditions used for growth assays in the presence of serum and treated overnight with ADT-007 (**A**), or serum starved and treated overnight with ADT-007 (**B**), or serum starved, treated overnight with ADT-007, and then stimulated for 10 min with EGF (**C**). Total RAS and GAPDH used as loading controls.

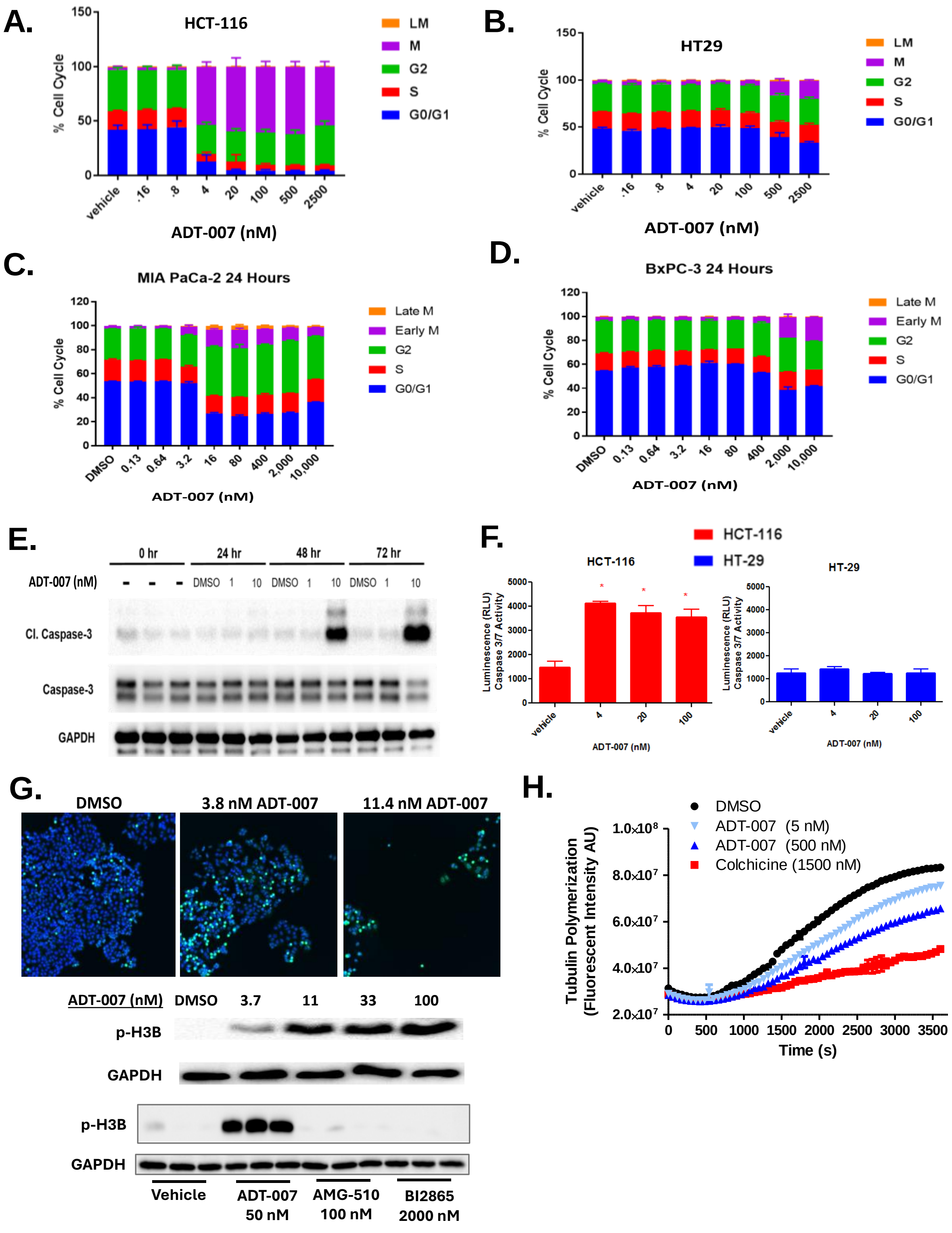


### Supplemental Figure 3

**Supplemental Figure 3. ADT-007 induces cell cycle arrest at G2/M phases and induces apoptosis of human CRC and PDA cell lines *in vitro*.** (**A-D**) ADT-007 induced G2/M cell cycle arrest in $KRAS^{G13D}$ HCT-116 (**A**) or $KRAS^{G12C}$ MIA PaCa-2 (**C**) cells, but not in $KRAS^{WT}$ HT-29 (**B**) or BxPC-3 (**D**) cells following 24 h of treatment. Error bars represent SEM of quadruplicate samples. (**E**) Time-dependent induction of apoptosis in $KRAS^{G12C}$ mutant MIA PaCa-2 cells by ADT-007 as measured by cleaved caspase-3 WB. (**F**) ADT-007 increases cleaved-caspase-3/7 levels in $KRAS^{G13D}$ HCT-116 cells but not in $KRAS^{WT}$ HT-29 as measured by a luminescence assay (ProMega). (**G**) ADT-007 increased histone 3B in association with reduced cell number as determined by immunofluorescence microscopy (top panel). ADT-007, but not AMG-510 or BI-2865, increased histone 3B, as determined by WB (bottom panel). (**H**) Kinetic assay of polymerization of fluorescently labeled, purified porcine tubulin incubated with indicated duplicate samples of ADT-007 for 1 h. Colchicine was included as a reference inhibitor of tubulin polymerization. Error bars represent SEM.

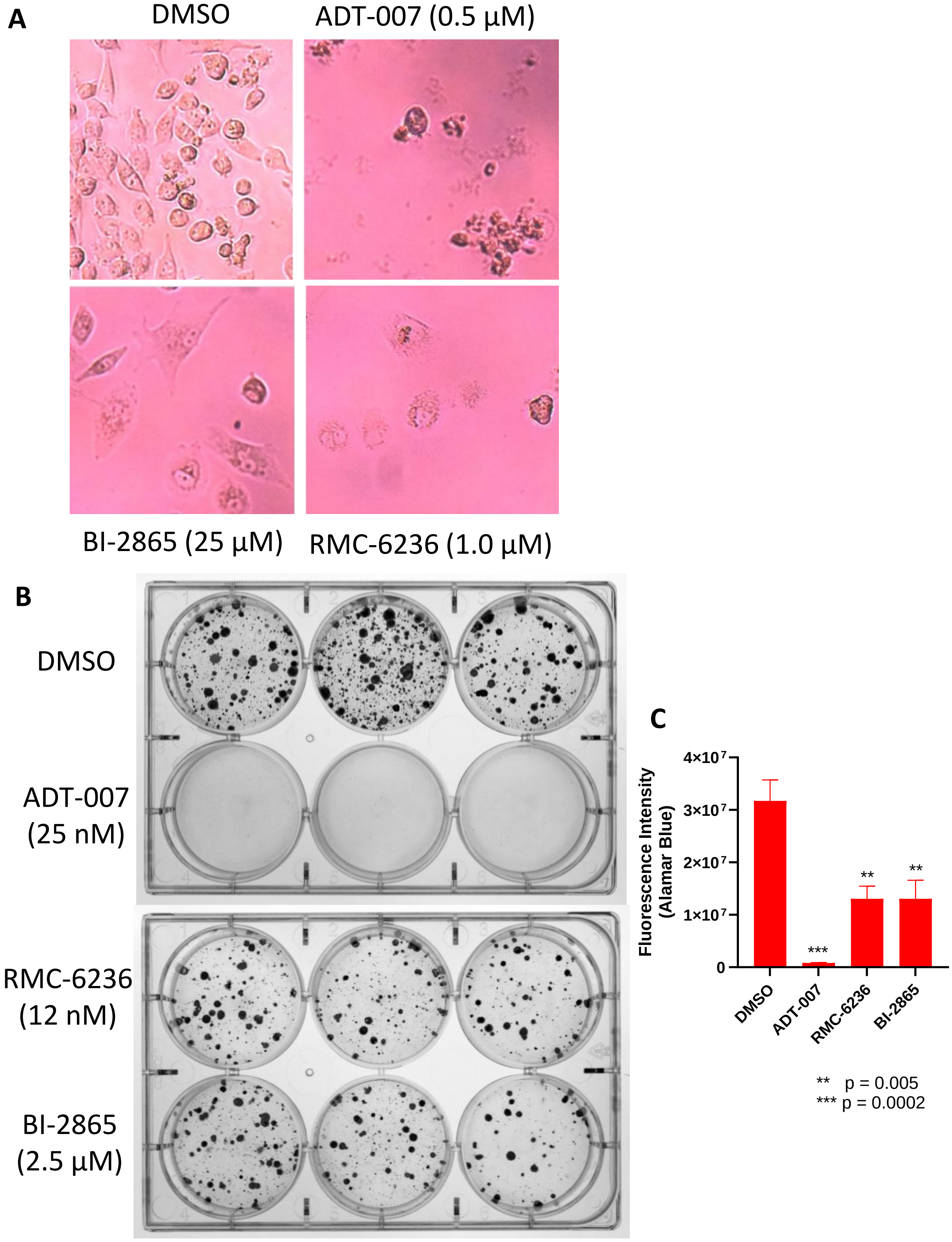


**Supplemental Figure 4**

**Supplemental Figure 4. ADT-007 is cytotoxic; BI-2865 (pan-KRAS inhibitor) and RMC-6236 (pan-RAS inhibitor) are cytostatic.** (**A**) Photomicrographs of ADT-007, BI-2865, and RMC-6236 treated $KRAS^{G12C}$ MIA PaCa-2 cells after 96-h treatment showing rounded condensed cells from ADT-007 treatment but attached and apparently viable cells from BI-2865 and RMC-6236 treatment. (**B-C**) Inhibition of MIA PaCa-2 cell colony formation (B) after 12-days of growth following a 48-h treatment period with ADT-007, BI-2865, or RMC-6236 showing near complete inhibition by ADT-007 while BI-2865 and RMC-6236 were appreciably less effective. Relative growth inhibition (C) was measured using the fluorescent Presto Blue HT (alamar blue) assay. Error bars represent mean ± SEM of triplicates. One-way ANOVA; ***p*=0.005, ***p=0.0002 *vs*. vehicle (DMSO) control.

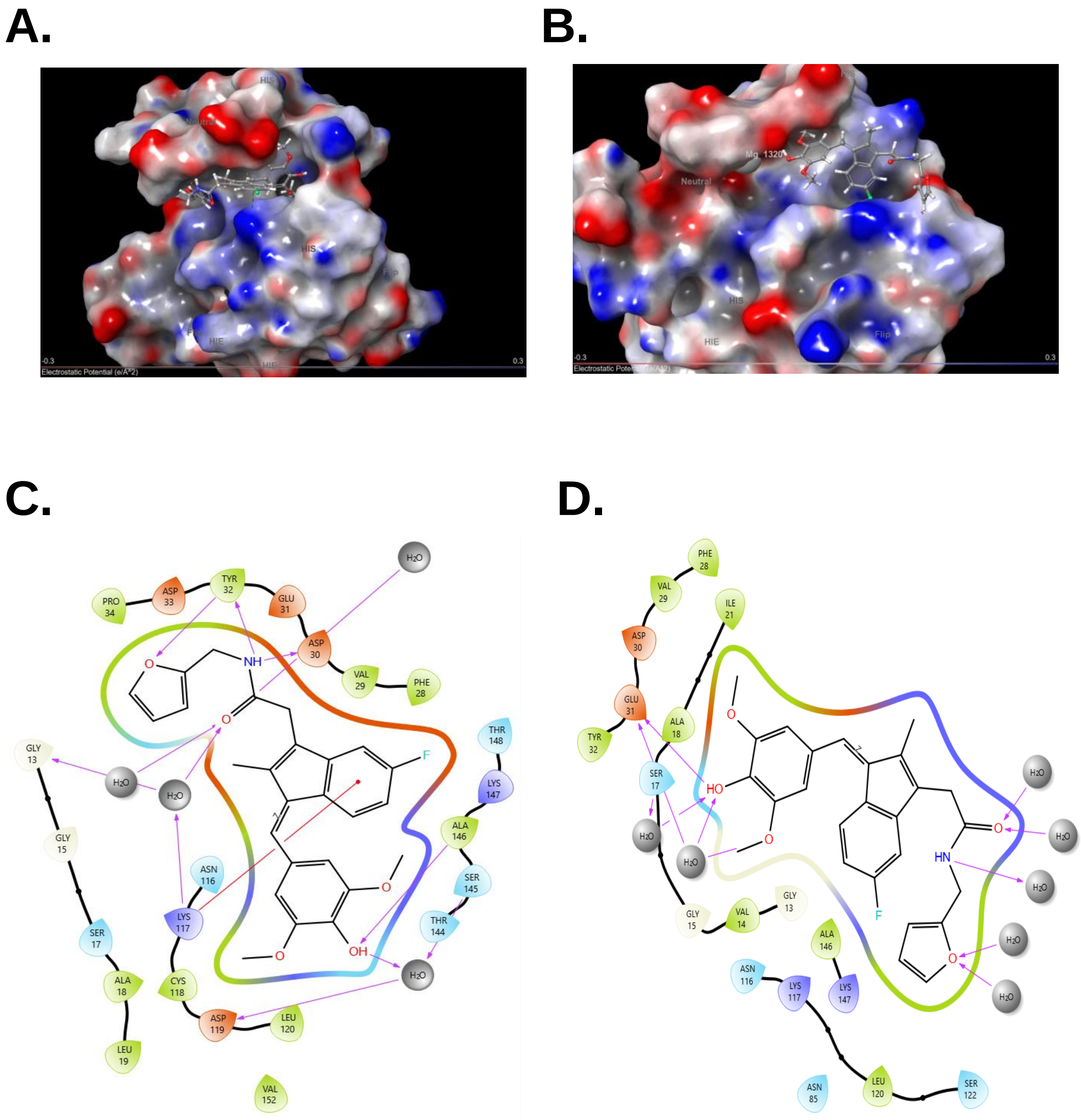


**Supplemental Figure 5**

**Supplemental Figure 5. Molecular modeling of ADT-007 binding to $KRAS^{WT}$.**

**(A)** Docking of ADT-007 to 4OBE ($KRAS^{WT}$ $Mg^{2+}$ absent). Docking Score -12.06. **(B)** Docking of ADT-007 to 4OBE ($KRAS^{WT}$ $Mg^{2+}$ present). Docking Score -11.08. In (**A**) and (**B**) 4OBE is rendered as a surface representation with the electrostatic potential mapped onto the surface. ADT-007 is rendered as a ball-and-stick representation. **(C)** Interaction diagram of docking of ADT-007 to 4OBE ($KRAS^{WT}$ $Mg^{2+}$ absent). Docking Score -12.06. **(D)** Interaction diagram of docking of ADT-007 to 4OBE ($KRAS^{WT}$ $Mg^{2+}$ present). Docking Score -11.08.

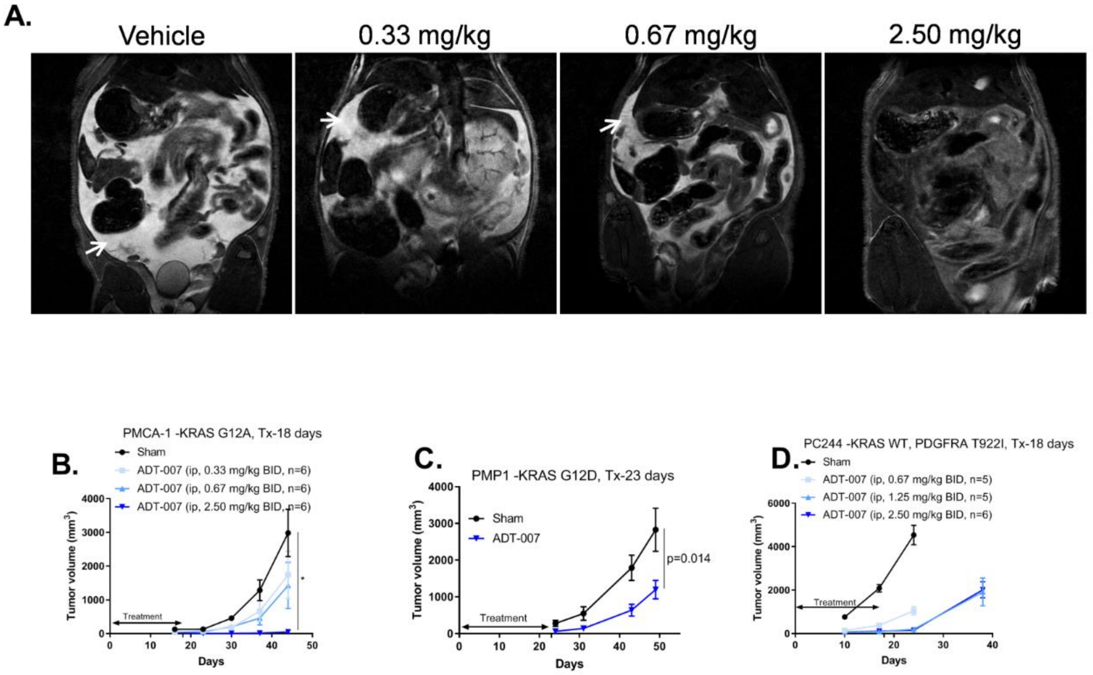


**Supplemental Figure 6**

**Supplemental Figure 6. ADT-007 inhibits growth of peritoneal metastases in mucinous CRC PDX in a dose dependent fashion.** PDX CRC cells were implanted IP in athymic mice. Antitumor activity of ADT-007 was measured by mucin volume using MRI**.** (**A**) Representative MR images of mice treated with vehicle, 0.33, 0.67 or 2.50 mg/kg ADT-007 IP twice daily on days 0-17 taken on day 44 after PMCA-1 tumor cell implantation. White arrows indicate mucin. Effect of ADT-007 treatment on tumor growth of (**B**) PMCA-1 ($KRAS^{G12A}$), (**C**) PMP1 ($KRAS^{G12D}$), and (**D**) PC244 ($KRAS^{WT}$) CRC tumors. Tumor growth was monitored and quantified via MRI. Data are summarized as mean values and error bars indicate SEM and contains 5-6 mice/group**.**

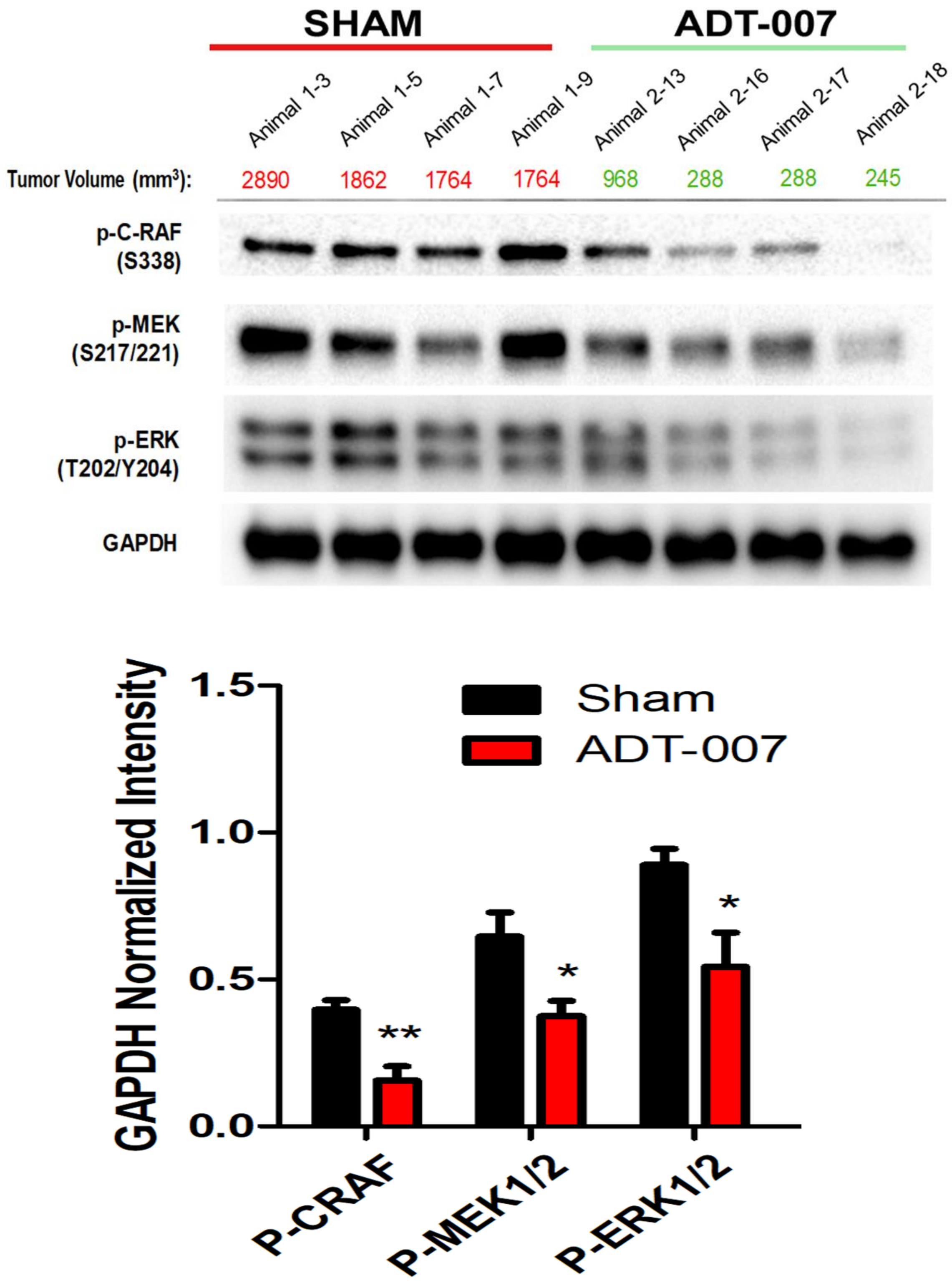


**Supplemental Figure 7**

**Supplemental Figure 7. ADT-007 inhibits tumor growth and MAPK signaling in the CT26 CRC mouse tumor model.** On the last day of ADT-007 treatment in $KRAS^{G12D}$ mutant CT26 CRC model, tumor lysates were prepared to determine the effect of ADT-007 on the RAS-MAPK pathway *in vivo*. (**top**) ADT-007 reduction of MAPK signaling in excised tumors from mice shown in **Figure 5E** after the last ADT-007 injection. Tumors were collected and homogenized to evaluate phosphorylated CRAF, MEK, and ERK by WB analysis. Each lane corresponds to a single mouse from the control (sham) or treatment groups (n=4). GAPDH was used as a loading control. Tumor weight for each mouse is listed above each lane. (**bottom**) Band intensity was quantified by ImageJ, normalized to GAPDH, and plotted as mean values with error bars indicating SEM (4 mice/group). * = $p<0.05$, ** = $p<0.01$.

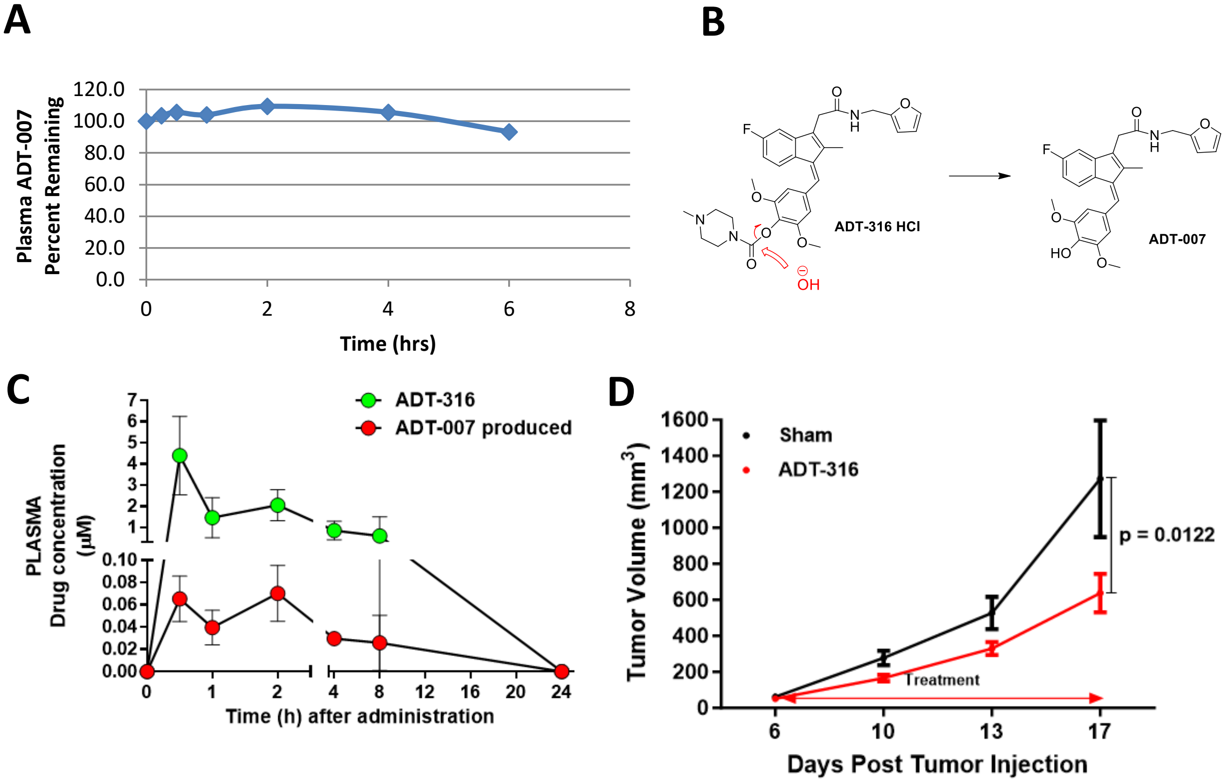


Supplemental Figure 8

**Supplemental Figure 8. ADT-007 prodrug, ADT-316, inhibits tumor growth.** (**A**) ADT-007 mouse plasma stability over time. (**B**) Chemical structure of ADT-316, a carbamate prodrug of ADT-007, and metabolism to ADT-007. (**C**) Pharmacokinetic profile of ADT-316 and generation of ADT-007 in female C57BL/6 mice following oral administration of ADT-316 (100 mg/kg) (mean ± SD, n=3-4 per time point). (**D**) ADT-316 inhibits growth of CT26 colon tumors in female BALB/c mice by oral administration of 100 mg/kg, BID for 12 days (mean ± SEM, n=8). Statistical significance was assessed using 2-way ANOVA.

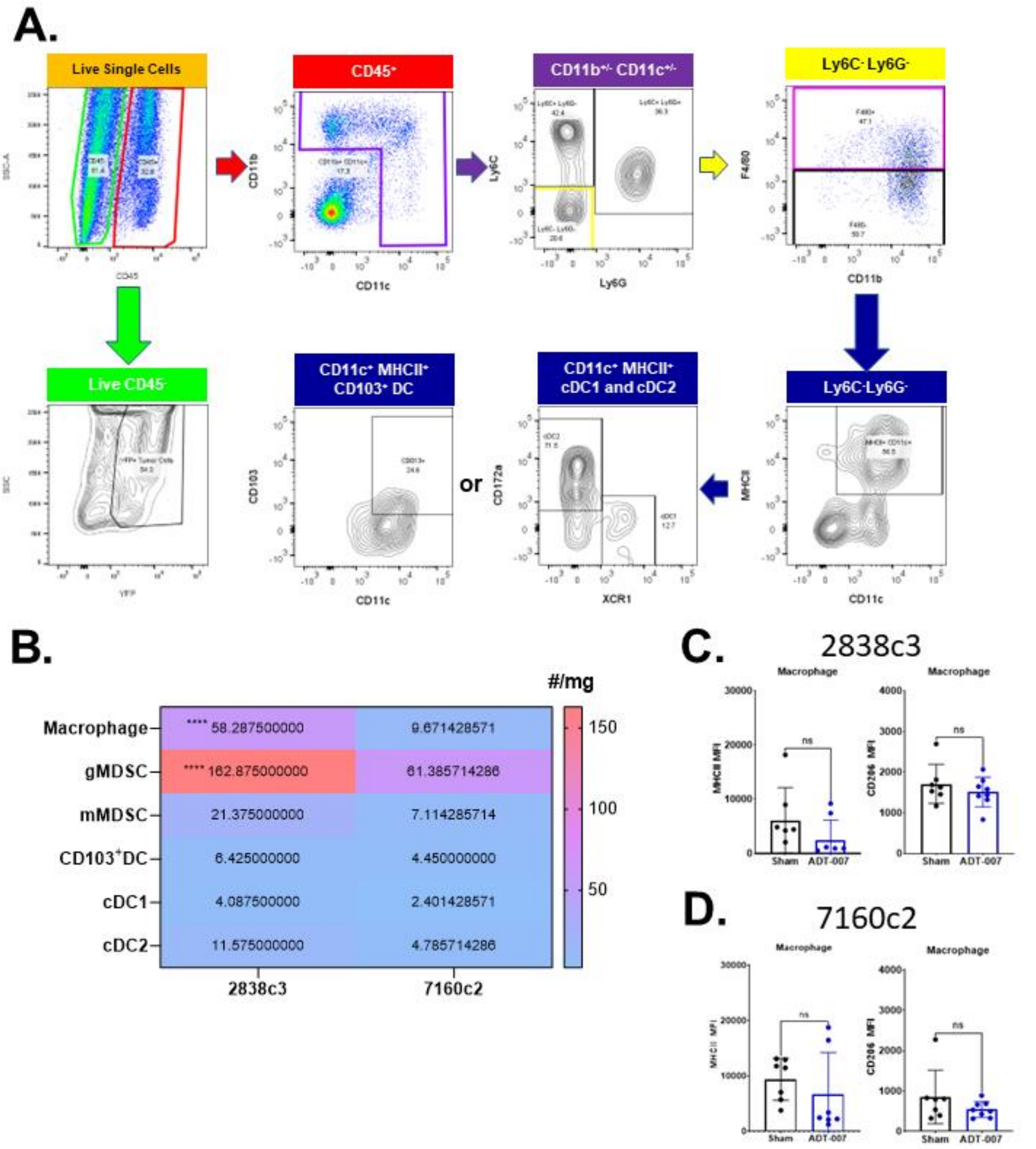


**Supplemental Figure 9**

**Supplemental Figure 9. Representative gating scheme for myeloid and tumor cells.** (**A**) Live-Dead Aqua negative $CD45^+$ and $CD45^-$ cells were gated upon and then myeloid cells were identified based on CD11b and CD11c expression, whereas tumor cells were identified by YFP fluorescence. The myeloid subsets were identified as follows: (i) mMDSC ($CD11b^+$ $Ly6C^{hi}$ $Ly6G^-$), gMDSC ($CD11b^+$ $Ly6C^{int}$ $Ly6G^+$), cDC2 ($CD11b^+$ $CD11c^+$ $MHCII^+$ $F4/80^-$ $CD172\alpha^+$), macrophage ($CD11b^+$ $Ly6C^-$ $Ly6G^-$ $F4/80^+$), and cDC1 (**$CD11b^-$ gate**: $CD11c^+$ $MHCII^+$ $XCR1^+$). (**B**) The total abundances (#/mg of tumor) of myeloid cell subsets were compared between sham-treated 2838c3 and 7160c2 mouse PDA tumors to determine differences in immune cell composition between the tumor models. Data were expressed as a heat map where mean values were obtained from n = 7-8 mice/group. Statistical significance was calculated using 2-way ANOVA and Tukey's correction for multiple comparisons. **** p <0.0001. (**C-D**) Expression (MFI) of MHCII and CD206 were evaluated in macrophages to determine activation and polarization status of specific subsets in (**C**) 2838c3 and (**D**) 7160c2 TiME. Data are representative of 2 independent experiments. p values were determined by Welch's t-test, ns indicates not significant. Error bars indicate SD.

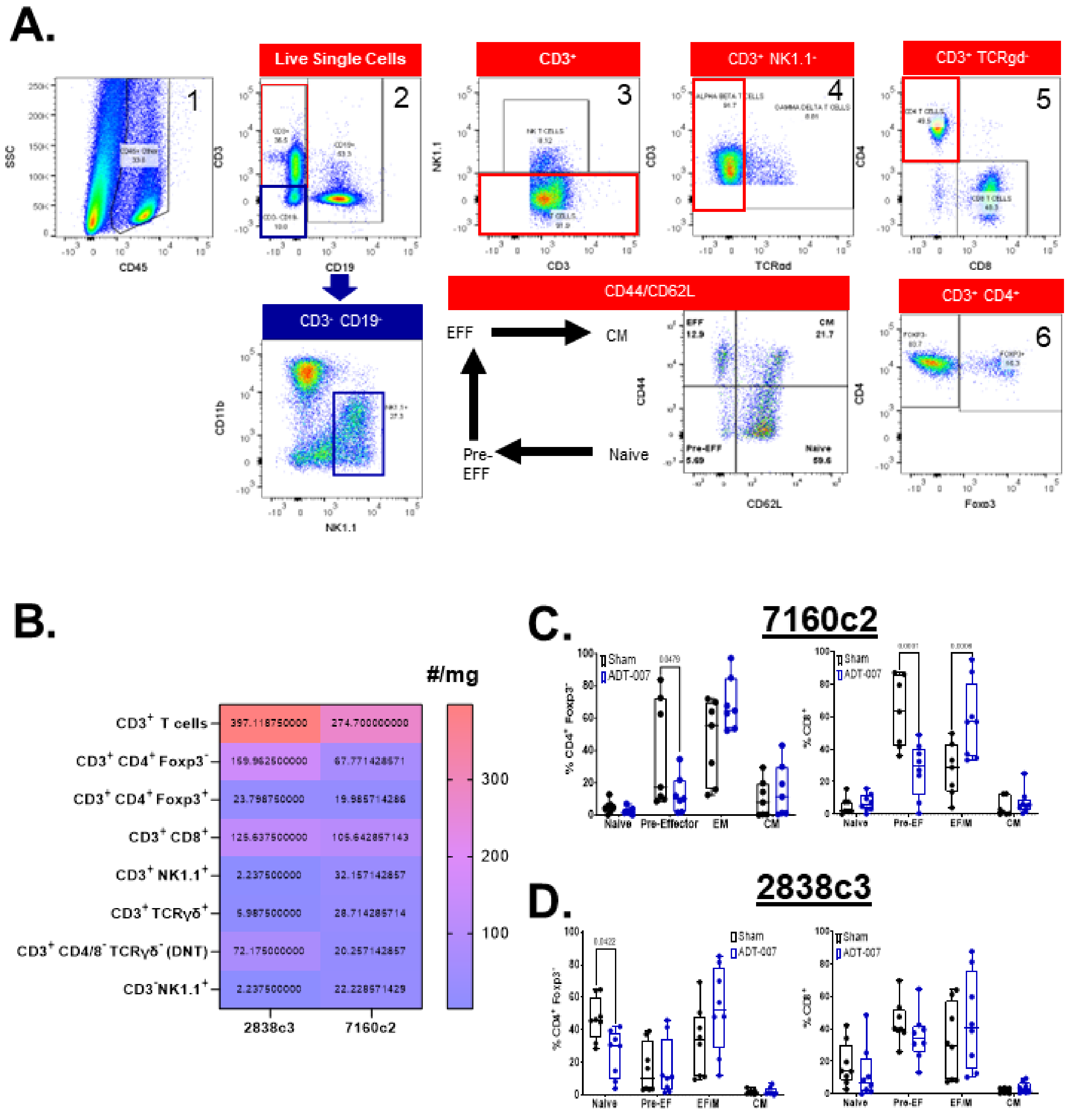


**Supplemental Figure 10**

**Supplemental Figure 10. Representative gating scheme for T and NK cells. (A)** Live-Dead Aqua negative $CD45^+$ cells were gated upon to identify NK and T cells as follows: NK ($NK1.1^+$ $CD3^-$), NKT ($NK1.1^+$ $CD3^+$), γδ T ($CD3^+$ $TCR\gamma\delta^+$), helper T ($CD3^+$ $CD4^+$) and cytotoxic T ($CD3^+$ $CD8^+$) T cells. Foxp3 expression within the $CD4^+$ T cell subset was used to identify $Foxp3^+$ regulatory T cells. Expression of PD-1, CD44, and CD62L were used to further identify activation status. (**B**) The total abundances (#/mg of tumor) of T cell and NK cell subsets were evaluated by comparing sham-treated 2838c3 and 7160c2 mouse PDA tumors (n = 7-8 mice/group) to determine differences in T cell composition in the TiME between each cell line. Data were expressed as a heat map where mean values were obtained from n = 7-8 mice/group. Statistical significance was calculated using 2-way ANOVA and Tukey's correction for multiple comparisons. (**C-D**) Expression of CD44 and CD62L in (**C**) 7160c2 and (**D**) 2838c3 tumor infiltrating $CD4^+$ and $CD8^+$ T cell subsets were evaluated by multi-parameter flow cytometry to determine percentages of naïve ($CD44^-$ $CD62L^+$), pre-effector ($CD44^-$ $CD62L^-$), effector memory ($CD44^+$ $CD62L^-$), and central memory T cells ($CD44^+$ $CD62L^+$) in ADT-007 and sham treated mice (n=8). Data are representative of 2 independent experiments. p values were determined by Welch's t-test. Error bars indicate SD.

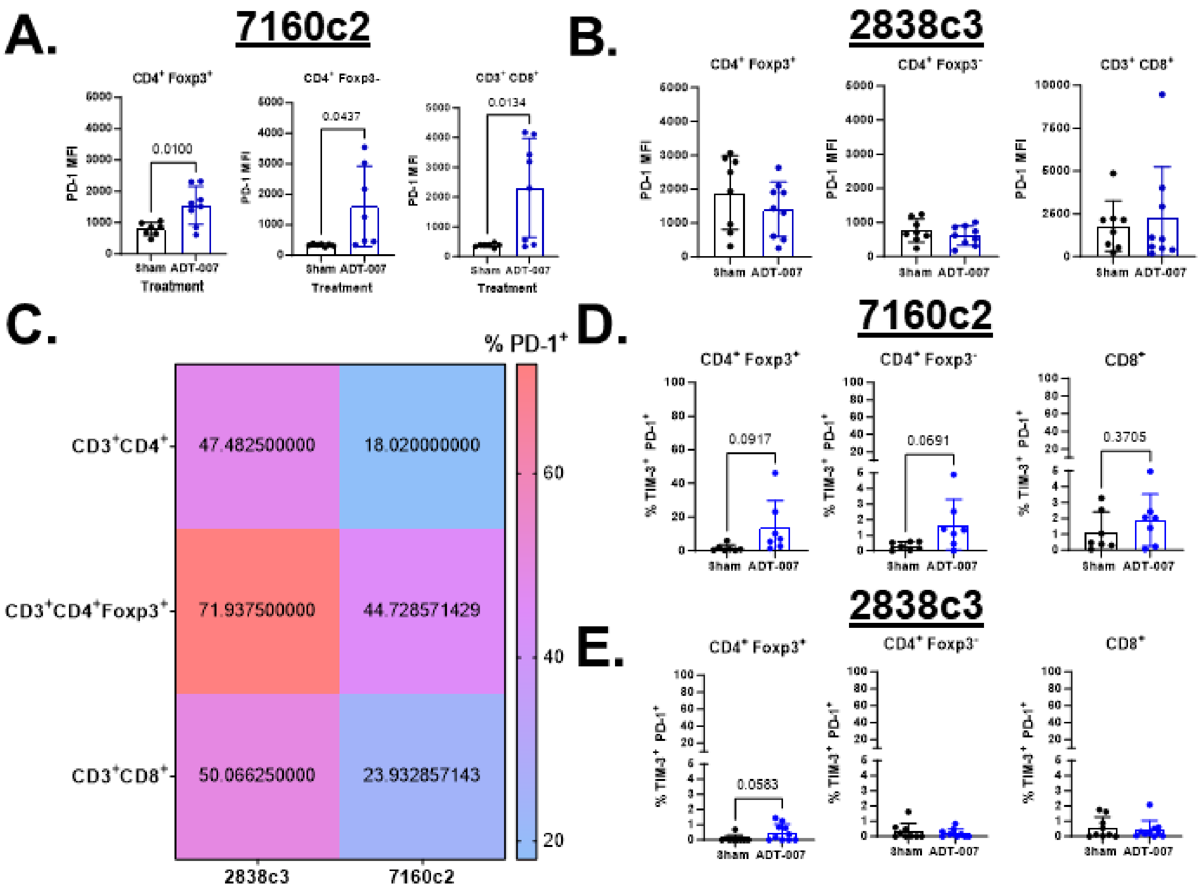


**Supplemental Figure 11**

**Supplemental Figure 11. Activation state and function of T cells in the 2838c3 and 7160c2 TiME.** **(A-B)** Expression of PD-1 in $CD4^+$ and $CD8^+$ T cell subsets from (**A**) 7160c2 and (**B**) 2838c3 TiME was graphed as mean fluorescent intensity. Data are representative of 2 independent experiments. p values were determined by Welch's t-test. Error bars indicate SD. (**C**) The expression levels of PD-1 on tumor infiltrating T cell subsets in sham treated 2838c3 and 7160c2 PDA tumor bearing mice were compared to determine activation state. T cell subsets from 2838c3 with exception of $CD3^+$ $NK1.1^+$ T cells expressed higher levels of PD-1 in sham treated tumors. Data were expressed as a heat map where mean values were obtained from n = 7-8 mice/group. Statistical significance was calculated using 2-way ANOVA and Tukey's correction for multiple comparisons. (**D-E**) Co-expression of PD-1 and TIM-3 on tumor infiltrating CD4 and CD8 T cells isolated from (**D**) 7160c2 and (**E**) 2838c3 tumor bearing mice treated with sham or ADT-007 (5 mg/kg, BID, SQ). Data are representative of 2 independent experiments. p values were determined by Welch's t-test. Error bars indicate SD.

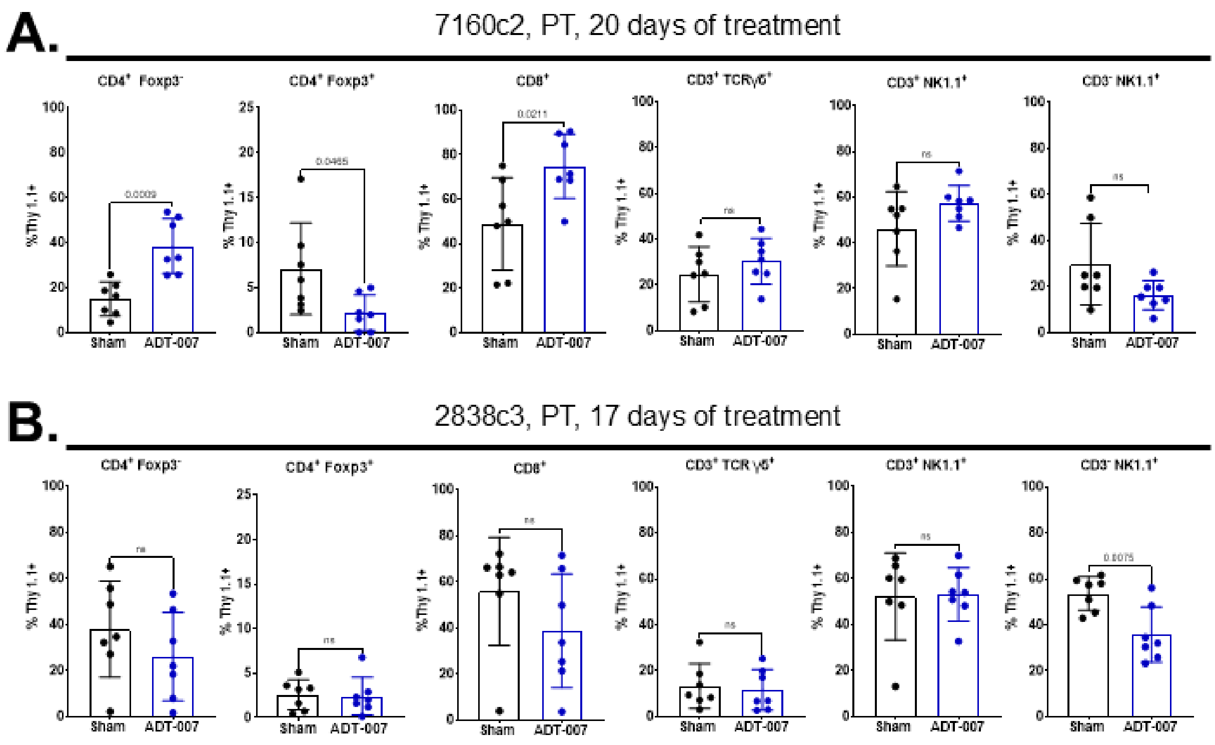


**Supplemental Figure 12**

**Supplemental Figure 12. ADT-007 treatment increases IFNγ production by $CD4^+$ and $CD8^+$ T cells in the 7160c2 TiME.** Production of IFNγ in T and NK cell subsets was evaluated *in vivo* by flow cytometry using Thy1.1-IFNγ reporter mice implanted with (**A**) 7160c2 or (**B**) 2838c3 mouse PDA tumor cells. Mice were treated with peritumoral injections of ADT-007 (5 mg/kg, BID, n = 7 mice/group) or sham for 21 days (7160c2) or 17 days (2838c3). P values were determined by Student's unpaired t-test, ns indicates not significant. Error bars indicate SD. Data are from a single experiment.

| Antibody | Supplier | Clone | Dilution | Reactivity/Use | Cat # |
|---|---|---|---|---|---|
| Total RAS | ThermoFisher | Ras10 | 1-1000 | Mouse/Human | MA1-012 |
| KRAS | Santa Cruz | F234 | 1-500 | Mouse/Rat/ Human | SC-30 |
| HRAS | Santa Cruz | 259 | 1-500 | Mouse/Rat/ Human | SC-35 |
| NRAS | Santa Cruz | F155 | 1-250 | Mouse/Rat/ Human | SC-31 |
| Total RAF | Cell Signaling | Polyclonal | 1-1000 | Human/ Mouse/ Rat/ Monkey | 9422 |
| p-C-RAF (S338) | Cell Signaling | 56A6 | 1-1000 | Mouse/Human Rat/Monkey | 9427 |
| Total MEK1/2 | Cell Signaling | Polyclonal | 1-1000 | Mouse/Human Rat/Monkey | 9122 |
| p-MEK1/2 (S217/221) | Cell Signaling | 41G9 | 1-1000 | Mouse/Human | 3958 |
| Total ERK1/2 | Cell Signaling | 137F5 | 1-1000 | Mouse/Human | 4695 |
| p-ERK1/2 (T202/Y204) | Cell Signaling | D13.14.4E | 1-2000 | Mouse/Human | 4370 |
| Total AKT | Cell Signaling | C67E7 | 1-1000 | Mouse/Human | 4691 |
| p-AKT (Ser473) | Cell Signaling | D9E | 1-1000 | Mouse/Human | 4060 |
| p-AKT (Thr308) | Cell Signaling | D25E6 | 1-1000 | Mouse/Human | 13038S |
| GAPDH | Cell Signaling | D16H11 | 1-1000 | Mouse/Human | 5174S |

**Supplemental Table 1. List of antibodies used for Western blot assays.**

| Antibody | Supplier | Clone | Dilution | Cat# |
|---|---|---|---|---|
| PD-1 FITC | BioLegend | 29F.1A12 | 1-200 | 135214 |
| CD206 FITC | BioLegend | C068C2 | 1-200 | 141703 |
| CD107α | BioLegend | ID4B | 1-1000 | 121607 |
| TIM-3 PE | BioLegend | B8.2C12 | 1-200 | 134003 |
| F4/80 PE | BioLegend | BM8 | 1-800 | 123110 |
| IL-17F PE | BioLegend | 9D3.1C8 | 1-800 | 517008 |
| Thy1.1 PE | BD | OX-7 | 1-200 | 551401 |
| CD19 PE Cy7 | BioLegend | ID3 | 1-800 | 152418 |
| Granzyme B PE Cy7 | BioLegend | GB11 | 5 uL/$10^6$ cells | 372214 |
| Ly6C PE Cy7 | BioLegend | HK1.4 | 1-3300 | 128017 |
| CD11b PE Dazzle | BioLegend | M1/70 | 1-3300 | 101255 |
| IFNγ PE Dazzle | BioLegend | XMG1.2 | 1-1000 | 505846 |
| CD45 PerCP | BioLegend | 30-F11 | 1-800 | 103130 |
| Foxp3 APC | Invitrogen | FJK-16s | 1-200 | 17-5773-82 |
| CD11c APC | BioLegend | N418 | 1-400 | 117309 |
| TNFα APC | BioLegend | MP6-XT22 | 1-1000 | 506308 |
| CTLA-4 APC R700 | BD | UC10-4F10-11 | 1-400 | 565778 |
| CD172α Alexa Flour 700 | BioLegend | P84 | 1-1000 | 144022 |
| CD62L APC Cy7 | BioLegend | MEL-14 | 1-800 | 104428 |
| CD86 APC Cy7 | BioLegend | GL1 | 1-200 | 105045 |
| IL-2 APC 750 | BioLegend | JES6-5H4 | 1-1000 | 503832 |
| CD3 BV421 | BioLegend | 145-2C11 | 1-800 | 100341 |
| XCR1 BV421 | BioLegend | ZET | 1-1000 | 148216 |
| Live/Dead Aqua | Invitrogen | n/a | 1-500 | L34966 |
| Ly6G BV570 | BioLegend | 1A8 | 5 uL/$10^6$ cells | 127629 |
| TCR γδ BV 605 | BioLegend | GL3 | 1-800 | 118219 |
| PD-L1 BV 605 | BioLegend | MIH5 | 1-400 | 153606 |
| CD4 BV 650 | BioLegend | GK1.5 | 1-800 | 100469 |
| MHCII BV 650 | BioLegend | M5/114/15.2 | 1-800 | 107641 |

| | | | | |
|---|---|---|---|---|
| NK1.1 BV 711 | BioLegend | PK136 | 1-400 | 108475 |
| CD103 BV 711 | BioLegend | 2E7 | 1-800 | 121435 |
| CD8 BV 785 | BioLegend | 53-6.7 | 1-800 | 100750 |
| CCR7 BV785 | BioLegend | 4B12 | 1-250 | 120217 |
| CD44 BUV 737 | BD | IM7 | 1-800 | 612799 |
| CD69 BUV 395 | BD | H1.2F3 | 1-400 | 569367 |
| B220 BUV 395 | BD | RA3.6B2 | 1-400 | 563793 |
| Fc Block (CD16/CD32) | BioLegend | 93 | 1-500 | 101320 |

**Supplemental Table 2. List of antibodies used for flow cytometry.**

| Type | Tissue Origin | Cell Line | Mutation Status | $IC_{50}$ (nM) | UGT1A1 | UGT1A6 | UGT1A10 | UGT2A1 | UGT2B10 | UGT2B15 |
|---|---|---|---|---|---|---|---|---|---|---|
| Mutant RAS, sensitive | Pancreatic | MiaPaCa2 | KRAS G12C | 2.1 | 0 | 0.287 | 0.263 | 0 | 0 | 0 |
| | Pancreatic | Panc1 | KRAS G12D | 2.4 | 0 | 0.057 | 0.043 | 0 | 0 | 0.014 |
| | Pancreatic | SW1990 | KRAS G12D | 6.3 | 0 | 0 | 0.138 | 0 | 0 | 0 |
| | Lung | A549 | KRAS G12S | 6.8 | 0.669 | 0.791 | 0.070 | 0 | 0.014 | 0.070 |
| | Pancreatic | CFPAC1 | KRAS G12V | 5.8 | 1.05 | 1.69 | 2.63 | 0 | 0 | 0.098 |
| | Colon | SW480 | KRAS G12V | 6.5 | 1.57 | 3.17 | 0.58 | 0 | 0 | 0 |
| | Breast | MDA231 | KRAS G13D | 3.6 | 0.214 | 0.807 | 0.043 | 0 | 0 | 0 |
| | Colon | HCT116 | KRAS G13D | 4.7 | 0.029 | 0.189 | 0 | 0 | 0 | 0 |
| | Ovarian | OVCAR8 | KRAS P121H | 5.8 | 0 | 0.214 | 0.057 | 0 | 0 | 0 |
| | Breast | HS578 | HRAS G12D | 3.9 | 0.043 | 0.163 | 0.098 | 0 | 0 | 0 |
| | Skin | SKMEL2 | NRAS Q61K | 2.1 | 0 | 0 | 0 | 0 | 0 | 0.084 |
| | Lung | H1299 | NRAS Q61K | 2.4 | 0 | 0.043 | 0 | 0 | 0 | 0 |
| WT RAS, resistant | Normal Lung | NHAEC | WT RAS | 19100 | * | * | * | * | * | * |
| | Normal Colon | NCM460 | WT RAS | 50000 | * | * | * | * | * | * |
| Downstream mutation (WT RAS) resistant | Ovarian | OV90 | WT RAS/mut BRAF | 350 | 5.00 | 6.92 | 4.57 | 0.070 | 0.057 | 0.632 |
| | Colon | HT29 | WT RAS/mut BRAF | 512 | 3.55 | 5.53 | 6.97 | 0 | 0.214 | 0.660 |
| | Pancreatic | BXPC3 | WT RAS/mut BRAF | 2500 | 2.11 | 6.41 | 7.75 | 0.057 | 0 | 0 |
| Upstream RAS activation sensitive | Lung | H1975 | WT RAS/mut EGFR | 3.9 | * | * | * | * | * | * |
| | Skin (mouse) | B16 | WT RAS/mut PDGFR | 5.8 | * | * | * | * | * | * |
| | Brain | U87MG | WT RAS/mut NF1 | 1.6 | 0 | 0.287 | 0 | 0 | 0 | 0 |
| | Ovarian | SKOV3 | WT RAS/mut NF1 | 6.7 | 2.89 | 7.66 | 0.202 | 0 | 0 | 0 |
| Amplified WT RAS, sensitive | Gastric | MKN-1 | Amplified WT KRAS | 4.4 | 0.95 | 4.12 | 0.490 | 0.09 | 0 | 0 |

**Supplemental Table 3. Sensitivity of cancer cells to ADT-007 is associated with high activated RAS levels (from H/N/K RAS mutations or upstream activators) and low UGT isozyme mRNA levels.** Growth inhibitory ($IC_{50}$) values were determined by CTG assay following 72-h of treatment. TPM normalized expression of the most significant differentially expressed UGT enzymes were obtained from data published in the Cancer Cell Line Encyclopedia accessed *via* the DepMap Portal hosted by the Broad Institute. * - expression data not available.

| Cell Line | Mutation Status | ADT-007 $IC_{50}$ (nM) | ADT-007 + propofol $IC_{50}$ (nM) | Fold Change |
|---|---|---|---|---|
| MIA PaCa-2 | KRAS G12C | 1.8 | 1.3 | 1.4 |
| HCT-116 | KRAS G13D | 7.3 | 2.4 | 3.0 |
| BxPC-3 | WT RAS/mut BRAF | 1813 | 98.3 | 18.4 |
| HT-29 | WT RAS/mut BRAF | 3160 | 122 | 25.9 |

**Supplemental Table 4. Propofol increases sensitivity of $RAS^{WT}$ cancer cell lines to ADT-007 without significantly affecting sensitivity of RAS mutant cell lines.** Cell lines were either treated with ADT-007 alone or co-treated with ADT-007 and propofol (200 µM). Growth inhibitory activity was determined after 72-h of treatment. $IC_{50}$ values are representative of triplicates from two independent experiments.

**Supplemental Methods**

High content image-based cell cycle analysis: Cells (5,000) were seeded per well in Perkin Elmer ViewPlate$^{TM}$ microplates (Perkin Elmer) and incubated overnight at 37°C. The following day, five-fold dilutions of ADT-007 or vehicle (DMSO) in dosing medium were added in equal volume (100 µL), to each well on the assay plate and incubated for 24 h. At the end of the treatment period, 150 µL per well of neutral buffered formalin fixative (10% paraformaldehyde) was rapidly added to each plate and incubated at room temperature for 20 min. Fixative was removed by washing with PBS. Plasma membranes were permeabilized by incubation with 0.3% Triton X-100 and 5% FBS in PBS for 1 h. TBS with 1 µg/mL DAPI was added to quantitate double stranded DNA within each cell nucleus. DNA content in each cell was determined by total DAPI stain intensity and assigned to the appropriate category (G0/G1, S, G2, M) using the MetaXpress cell cycle analysis program. Results from each treatment were reported as average cell number in each of nine fields, as well as the percentage of cells within each phase of the cell cycle. Each of these parameters was analyzed using GraphPad Prism, and, where appropriate, $IC_{50}$ values have been reported.

Spheroid Cell Culture: MIA PaCa-2 cells were seeded at a density of $10^3$ to 1.5 x $10^3$ cells into single wells of 96 well cell culture plates coated with a 1.5% agarose. Cells were cultured for 72 h, then 50% of the media was replaced with either media containing vehicle (DMSO at 1%) or increasing concentrations of ADT-007 (0.1, 0.5, 1, 5, 10, 50, 100, 1000 nM) for up to 10 days. Cells were then dissociated, and cell numbers were enumerated and $IC_{50}$ calculated.

Colony formation assays: Approximately 500 cells/well (6-well plate) were incubated for 24 h at 37°, dosed 24 h later with either increasing concentrations of ADT-007 (0.5. 5, 20, 30, 40, 50, 70 nM), or DMSO and then incubated for period of 8-15 days. Endpoints were determined when colonies in the vehicle control wells reached a size of approximately 50 cells or larger. At endpoint, wells were washed with PBS, then stained for 1 h at RT in 2 mL of crystal violet solution in 1% formalin and 1% methanol. Plates were imaged using a ChemiDoc Imaging System (Bio-Rad), and colonies were counted using a custom macro in Nikon Elements Research developed by Dr. Joel Andrews (University of South Alabama).

Fluorometric growth inhibition assay: In PrestoBlue HT (ThermoFisher) was added to culture medium (1/10 volume) of clonogenic assay plates and incubated for 30 min. 150 uL was transferred from each well to a 96-well assay plate and fluorescence (560ex/590em) was measured in a Molecular Devices ID5 multilabel plate reader. Relative fluorescence intensities were graphed using GraphPad Prism.

Fluorescent tubulin polymerization assay: A fluorescent tubulin polymerization assay kit (Cytoskeleton, Inc., Catalog #BK011P) was used according to the supplied protocol to determine effects of experimental compounds on tubulin polymerization. Briefly, control (vehicle) and test compounds (ADT-007, colchicine) were added to duplicate samples assay mix (100 ng porcine tubulin, 1 mM GTP, 25% glycerol, 2x assay buffer containing: 80 mM Piperazine-N,N'-bis[2-ethanesulfonic acid] sequisodium salt; 2.0 mM Magnesium chloride; 0.5 mM Ethylene glycol-bis(b-amino-ethyl ether) N,N,N',N'-tetra-acetic acid, pH 6.9, 10 μM fluorescent reporter( 4’, 6-Diamidino-2-phenylindole). The plate was immediately transferred to a Molecular Devices Spectramax Paradigm fluorescence plate reader equilibrated to 37°C. Fluorescence (360ex/450em) was read each 1 min for 60 min at 37°C.

Molecular modeling: $KRAS^{WT}$ (PDB entry 4OBE) was used for all modeling studies. The protein was prepared using the Protein Preparation workflow within Schrodinger 2022-3. For the $Mg^{2+}$ free form, $Mg^{2+}$ was deleted, and the resulting structure was then submitted to the workflow. Initial docking studies with ADT-007 were performed with GDP removed from KRAS using GLIDE. The highest scoring docking pose was submitted to molecular dynamics using DESMOND for 100 nsec, and the resulting trajectory file was analyzed by molecular clustering with 12 clusters resulting from each dynamics run. The highest scoring pose was then determined using GLIDE and this pose was then optimized using induced-fit docking.

Mouse plasma stability assay: Mouse plasma was spiked with ADT-007 at a concentration of 1 μM and incubated in a shaking water bath at 37°C. Samples were taken at 0, 0.25, 0.5, 1, 2, 4, and 6 h, quenched with acetonitrile and analyzed by LC-tandem mass spectrometry to measure the remaining amount of ADT-007.

Pharmacokinetic studies: Pathogen-free 6-8-week-old female C57BL/6 mice were purchased from Envigo (strain 044) and housed in the Biologic Research Laboratory at the University of South Alabama, College of Medicine. Mice were housed in sterilized filter-stopped cages in a room with controlled temperature (24 ± 2 °C), 30 – 70% humidity, and a 12:12 h light-dark cycle. Mice were fed pelleted Teklad global 18% protein rodent diet (Teklad 2018, Inotiv Co.) *ad libitum* and received filtered (by reverse osmosis) municipal water *ad libitum* throughout the study. PK experimental protocol was approved by the IACUC of the University of South Alabama. Mice were acclimated for 2 wks before the start of the study. ADT-316 was dissolved in sterile water at a concentration of 10 mg/mL to yield a dose of 100 mg/kg and administered once by oral gavage. Blood was drawn at 0.5, 1, 2, 4, 8, and 24 h and collected into $K_2$EDTA-microtainer tubes followed by separation of plasma. ADT-316 levels in plasma as well as ADT-007 levels generated following oral administration of ADT-316 were determined using reverse phase chromatography with MS/MS detection.
